# Distributed representations of chemosensory valence in a naïve vertebrate brain

**DOI:** 10.64898/2026.06.17.732810

**Authors:** Bethan Jenkins, Thomas Frank

## Abstract

Chemical cues guide essential behaviors by signaling food, danger, and social information. A major dimension of chemosensory processing is valence, which biases animals toward approach or avoidance, yet its brain-wide organization in vertebrates remains unclear. Here, we used larval zebrafish to map neuronal activity associated with behaviorally defined appetitive and aversive chemosensory cues. Free-swimming assays identified a chemically diverse stimulus panel that elicited robust approach or avoidance, and key valence-dependent motor signatures were preserved during partially immobilized two-photon Ca²⁺ imaging with simultaneous tail tracking. Brain-wide activity revealed partly distinct patterns of stimulus identity, valence-related activity, and movement-correlated neuronal recruitment. Stimulus identity was strongest in the olfactory bulb and pallium, whereas small, coordinated valence-related populations were distributed across multiple telencephalic and diencephalic regions. These populations were partly separable from movement-correlated neurons, especially in forebrain regions, but showed valence-specific, opposing relationships with motor output. Together, our findings show that chemosensory valence-related activity in a naïve vertebrate brain is not confined to a single sensory or motor-associated region, but is organized across a distributed, regionally structured brain-wide architecture.

**HIGHLIGHTS:** Naïve larval zebrafish show appetitive and aversive chemosensory behaviors that are preserved during imaging

Chemosensory stimuli evoke widespread brain activity, revealing distributed stimulus identity and valence representations

Valence-encoding neurons form small, coordinated populations across multiple forebrain regions

Valence-related activity is partly separable from movement-correlated activity

## INTRODUCTION

Animals depend on chemical cues to locate food, avoid danger, and guide social behavior ^1,2^. Many such cues carry innate appetitive or aversive behavioral significance, biasing animals toward approach or avoidance ^3^. Although chemosensory valence is central to odor-guided behavior, the brain-wide circuit architecture through which valence-related activity is represented and organized remains incompletely understood. Defining this architecture is essential for understanding how chemosensory information progresses from stimulus detection toward brain activity that can guide behavior.

Work across species supports the idea that chemosensory valence is shaped by distributed circuits rather than by early sensory representations alone. In insects, appetitive and aversive odors recruit olfactory, associative, and downstream motor-linked pathways, providing powerful circuit-level models for how odor value can guide behavior ^4–9^. In mammals, innately attractive or aversive odors, as well as learned odor values, have been associated with activity across the olfactory bulb ^10–13^, piriform cortex ^14,15^, amygdalar complex ^16–20^, olfactory tubercle ^21–25^, hypothalamic circuits ^26–29^, and frontal and orbitofrontal regions ^17,30^. In zebrafish, an olfactory cortex homolog maps odor space onto a learning-modified valence representation ^31^ and chemosensory cues can trigger specific neuronal activity patterns and sensorimotor programs ^32–34^. Although these studies differ substantially in species, methods, and behavioral context, they suggest that chemosensory valence is organized across multiple processing levels.

However, the organizational principles by which valence-related activity is distributed across the brain, and related to stimulus identity and motor output, remain largely unclear. Appetitive and aversive cues may recruit shared downstream circuits, partly distinct distributed populations, or region-specific combinations of both ^35,36^. It also remains unclear where valence-related activity diverges from stimulus identity representations ^31^, and whether it remains separable from motor-correlated activity once chemosensory cues begin to bias behavior. These questions are particularly important because approach and avoidance provide the operational readout of valence, yet the neural activity associated with valence need not be identical to the activity most directly coupled to movement.

Addressing these questions requires an experimental setting in which multiple appetitive and aversive cues can be tested under matched conditions, while neuronal activity and behavior are measured simultaneously across the brain. Such an approach allows stimulus identity, valence-related activity, and motor-correlated signals to be compared within the same animals and across brain regions, rather than inferred from separate assays or selected candidate areas.

Larval zebrafish are well suited for this purpose because their small, optically accessible brain enables cellular-resolution activity measurements across large parts of the nervous system during controlled sensory stimulation ^37,32–34^. At early developmental stages, larvae already show chemosensory behaviors, including responses to appetitive and aversive cues ^38–43,32,44,33,34^, while their behavioral repertoire remains comparatively simple. This provides an opportunity to define a behaviorally validated stimulus set and link brain-wide chemosensory activity to motor output in an early-life vertebrate circuit context. This stage also precedes extensive experience-dependent modification ^45,46^ and later expansion of associative circuitry ^47,48^, allowing valence-related activity to be studied before stronger learning-dependent influences emerge.

Here, we used larval zebrafish to map neuronal activity associated with behaviorally defined chemosensory valence across the brain. We first established a stimulus panel that elicited appetitive or aversive behavioral responses in freely swimming larvae, and then tested how key valence-dependent motor signatures were preserved during partially immobilized two-photon Ca²⁺ imaging with simultaneous tail tracking. This allowed us to compare stimulus identity, valence-related activity, and movement-correlated neuronal populations under matched stimulus conditions. We find that appetitive and aversive cues evoke widespread but regionally structured activity, with stimulus identity represented most strongly in the olfactory bulb and pallial regions and valence-related activity distributed across multiple telencephalic and diencephalic regions. Valence-related populations were partly separable from movement-correlated neurons, especially in anterior regions, while posterior diencephalic and hindbrain regions showed closer alignment with motor output. Together, these findings reveal a distributed brain-wide architecture for chemosensory valence-related activity already in a naive, early-life vertebrate brain.

## RESULTS

### Free-swimming screen identified stimuli with appetitive or aversive behavioral effects

To identify chemosensory stimuli that elicit approach or avoidance, we screened groups of freely swimming zebrafish larvae in an agarose patch assay adapted from established larval olfactory paradigms ^44^. Groups of approximately 20 larvae explored an arena containing a stimulus-loaded agarose patch on one side and a control patch on the other, allowing attraction or avoidance to be quantified from larval position relative to the stimulus source (**Fig. 1A,B**; **Methods**). Because the stimulus gradient developed gradually and concentrations decreased steeply with distance from the patch, behavioral analyses focused on the final five minutes of the ten-minute exposure (**Fig. S1A-D**). Each of the seven candidate stimuli was tested in more than 135 larvae across 8 to 12 arena recordings.

**Figure 1.**
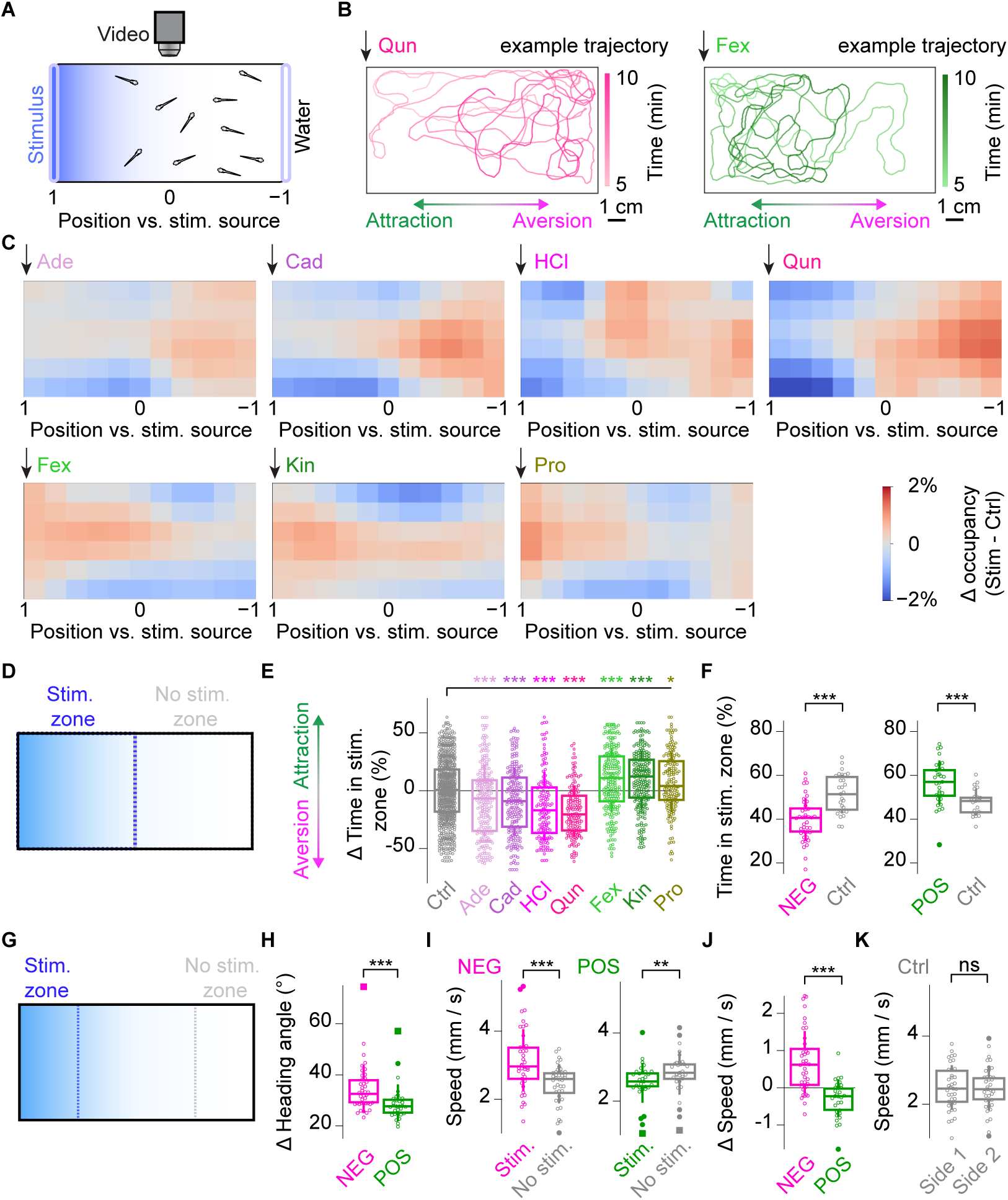
**Behavioral responses to chemosensory stimuli in freely swimming, naïve zebrafish larvae**. (**A**) Free-swimming assay. In a single experiment, ∼20 larvae explored an arena for 10 min while stimulus gradients formed from agarose patches. (**B**) Example single-larva trajectories showing avoidance of quinine (Qun; left) and attraction to food extract (Fex; right), both diffusing from the left side (arrow). Trajectories show the last 5 min of recording and are color-coded by time, with darker colors indicating later time points. (**C**) Heatmaps show differential occupancy (Δ occupancy; stimulus − no-stimulus control), calculated by normalizing occupancy within each spatial bin and subtracting the corresponding normalized occupancy from separate control experiments. Maps were aligned so that the stimulus source was always on the left (arrow). Blue: reduced occupancy; red: increased occupancy. (**D**) Schematic of stimulus zone. (**E**) Box plots show stimulus-zone occupancy changes during the final 5 min after subtraction of experiment-matched, agarose-only controls. Significance was assessed by nonparametric multiple comparisons versus control with adjusted p-values; *p < 0.05, ***p < 0.001. Sample sizes: Fex, n = 226; Kin, n = 233; Pro, n = 161; Ade, n = 245; Cad, n = 213; HCl, n = 136; Qun, n = 151; Ctrl, n = 722 larvae. (**F**) Stimulus-zone occupancy at the individual-experiment level, pooled for aversive (NEG) and appetitive (POS) stimuli and compared with experiment-matched controls. Wilcoxon-Mann-Whitney test: NEG, N = 40; Ctrl, N = 29; p = 3.1 × 10⁻⁶; POS, N = 31; Ctrl, N = 21; p = 0.00025. N, number of experiments. (**G**) Schematic showing a more narrowly defined stimulus zone (Stim.) and the corresponding control zone near the agarose-only patch (No stim.), used for analyses in H-K. (**H**) Change in heading angle between consecutive bouts for aversive (NEG) and appetitive (POS) stimuli; Wilcoxon-Mann-Whitney test, p = 0.00062; N as in F. (**I**) Swim speed within the stimulus zone compared with the corresponding no-stimulus zone for aversive stimuli (left; Wilcoxon signed-rank test, p = 5.8 × 10⁻⁶; N = 40) and appetitive stimuli (right; p = 0.0025; N = 31). (**J**) Experiment-wise swim-speed change relative to the no-stimulus zone for aversive and appetitive stimuli; Wilcoxon-Mann-Whitney test, p = 7.9 × 10⁻⁸; N as in F. (**K**) Same as I for agarose-only control experiments; Wilcoxon signed-rank test, p = 0.81; N = 37.

Averaged spatial occupancy maps revealed clear stimulus dependent biases (**Fig. 1C**). Adenosine (Ade), cadaverine (Cad), hydrochloric acid (HCl), and quinine (Qun) reduced occupancy of the stimulus zone, consistent with avoidance, whereas food extract (Fex), kin water (Kin), and L-proline (Pro) increased occupancy, consistent with attraction. Quantification of time spent in the stimulus zone confirmed these effects at the population level (**Fig. 1D,E**). Comparison to control trials with two stimulus-free artificial fish water (AFW) patches established that the biases were chemosensory in origin: aversive stimuli (“NEG”: Ade, Cad, HCl, Qun) decreased occupancy relative to controls (Ade, p = 0.00080; Cad, p = 0.00024; HCl, p = 8.0 × 10⁻⁷; Qun, p = 3.2 × 10⁻¹²), whereas appetitive stimuli (“POS”: Fex, Kin, Pro) increased it (Fex, p = 0.00016; Kin, p = 1.8 × 10⁻⁵; Pro, p = 0.015). Preference distributions were broad for all stimuli, indicating substantial larva to larva variability within each stimulus condition, yet cohorts differed significantly from controls in all cases. Because attraction and aversion were also evident at the level of individual arena experiments (**Fig. 1F**), experiment to experiment variability was unlikely to account for the majority of this heterogeneity.

We next asked whether attractive and aversive stimuli also differed in their effects on navigation near the stimulus patch (**Fig. 1G-K**). Aversive stimuli increased turning, as reflected by larger changes in heading angle between bouts (p = 0.00062; **Fig. 1H**). Relative to the non-stimulus side within the same experiments (**Fig. 1G**), aversive stimuli also increased swim speed (p = 5.8 × 10⁻⁶ ; **Fig. 1I**), whereas appetitive stimuli decreased it (p = 0.0025; **Fig. 1I**), yielding clear experiment-wise differences between the two valence classes (p = 0.0002; **Fig. 1J**). No side difference was detected in control experiments with two empty agarose patches (p = 0.81; **Fig. 1K**). Similar observations were made when the analysis was performed at the level of individual fish (**Fig. S1E-I**).

Thus, under the conditions tested here, aversive stimuli were associated with reduced stimulus-zone occupancy, increased turning, and higher swim speed, whereas appetitive stimuli were associated with increased occupancy and slower swimming. This identified four avoidance-associated stimuli (Ade, Cad, HCl, Qun) and three approach-associated stimuli (Fex, Kin, Pro) for subsequent testing under imaging-compatible conditions.

### Partially immobilized larvae retained key valence-dependent behavioral features

Here, we asked whether key behavioral differences between appetitive and aversive stimuli were preserved under conditions compatible with live imaging. We used a partially immobilized preparation that allowed controlled chemosensory stimulation, simultaneous tail tracking, and two-photon Ca²⁺ imaging, while leaving the nose, mouth, and tail free (**Fig. 2A**). Each fish was exposed to the full stimulus panel: three stimuli were presented at a single concentration, and four stimuli, two NEG (Cad, Qun) and two POS (Fex, Pro), were tested at three concentrations **(Methods)**. Stimulus concentration conditions were delivered repeatedly in randomized order within a continuous constant-flow artificial fish water (AFW) stream. Tail kinematics were tracked from high-speed infrared videos (**Fig. 2B,C**), swim bouts were identified from sliding-window changes in tail angle variability, referred to as tail vigor, and their counts were normalized to pre-stimulus levels (**Methods**).

**Figure 2.**
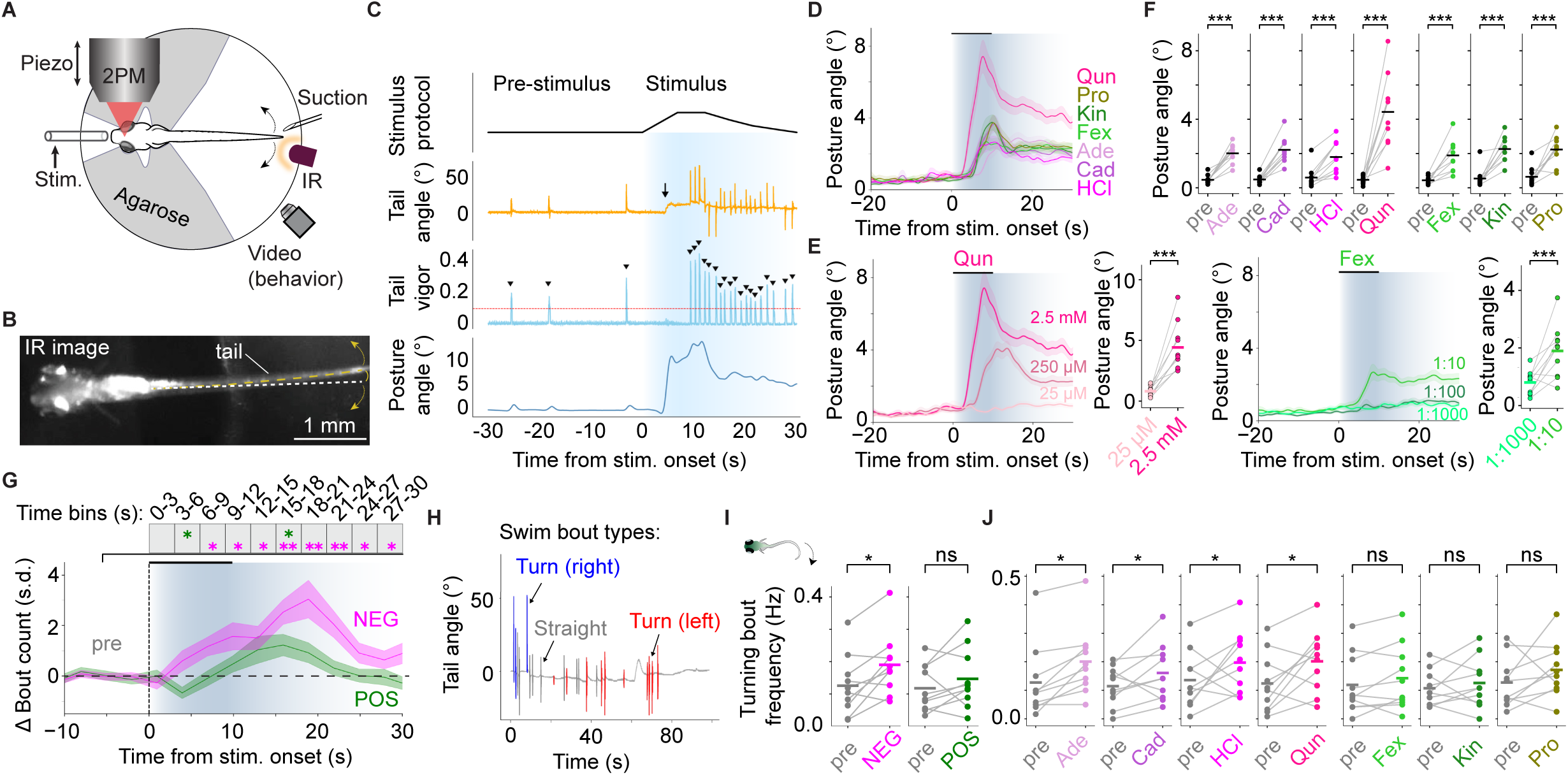
**Behavioral responses to chemosensory stimuli in partially immobilized, naïve zebrafish larvae**. (**A**) Partial-immobilization assay combining chemosensory stimulation, tail tracking, and volumetric two-photon microscopy (2PM) in agarose-embedded larvae with parts of the head and tail free. (**B**) Example infrared (IR) image used for tail tracking. (**C**) Example single-trial behavioral traces showing the stimulus’ temporal profile (top), raw tail angle (orange; arrow, postural-response onset), tail vigor (blue), detected swim bouts (black arrowheads), and bout-free tail posture angle after interpolation across bout periods (bottom). (**D**) Mean posture angle across all seven stimuli; highest concentration shown where applicable. Background shading indicates approximate stimulus profile; trace shading indicates s.e.m. (**E**) Mean posture angles for three concentrations of Qun (left) and Fex (right). For both stimuli, the highest concentration elicited greater posture angle responses than the lowest during the 20-s stimulus window; linear mixed-effects model with concentration as fixed effect, and fish identity (N = 10) and imaging session as random intercepts; ***p < 0.001. (**F**) Mean posture angle in 20-s pre-stimulus and stimulus windows. All stimuli, at the concentrations shown in D, elicited significant postural responses; mixed-effects model with time window (pre-stimulus vs. stimulus) as fixed effect; ***p < 0.001. (**G**) Change in total bout count over time, expressed as z-scored deviation from the pre-stimulus baseline and quantified in 3-s bins, averaged across highest-concentration appetitive (POS; green) and aversive (NEG; magenta) stimuli. Aversive stimuli increased bout counts after stimulus onset (dashed line) and remained elevated throughout the analysed window, whereas appetitive stimuli caused an initial suppression (3-6 s bin) followed by a delayed transient increase during washout (15-18 s bin); Wilcoxon signed-rank test versus baseline, *p < 0.05, **p < 0.01. Background shading indicates approximate stimulus profile; trace shading indicates s.e.m. (**H**) Example trace showing swim bouts classified as turns (blue/red) or straight bouts (gray), based on the mean tail angle during the first 70 ms of each bout (**Methods**). (**I**) Turning-bout frequency in 20-s pre-stimulus and stimulus windows. Aversive stimuli increased turning-bout frequency, whereas appetitive stimuli did not; Wilcoxon signed-rank test, NEG: p = 0.027, POS: p = 0.19. (**J**) Turning-bout frequency as in I for individual stimuli: Ade, 25 µM; Cad, 2.5 mM; HCl, pH 5; Qun, 2.5 mM; Fex, 1:10; Kin, 1:100; Pro, 2.5 mM; mixed-effects model as in F, *p < 0.05, ns, not significant.

In addition to discrete swim bouts, trials often showed stimulus-evoked, low-amplitude sustained tail deflections, with bouts superimposed, resembling previously described postural responses ^49,50^ (**Fig. 2C-F**). These posture angle responses were concentration dependent (highest vs. lowest concentration, p < 0.001; **Fig. 2E**; **Fig. S2A,B**) and robust for both appetitive and aversive stimuli at the highest concentrations (pre-stimulus vs. stimulus, p < 0.001; **Fig. 2D,F**), indicating that all seven cues elicited detectable motor responses. In a separate control configuration, angled stimulus delivery elicited postural bends toward the direction of stimulus flow (**Fig. S2C**), consistent with an orienting-like motor component under partial immobilization ^49,51^.

At the highest concentrations, POS stimuli caused a transient decrease in overall bout frequency shortly after stimulus onset (3 to 6 s bin, p = 0.02; **Fig. 2G**), whereas NEG stimuli induced a prolonged increase in bout frequency (p < 0.05 for eight consecutive time bins after the 3 to 6 s bin; **Fig. 2G**). During the decline in stimulus concentration (**Fig. S2D**), POS trials also showed a brief increase in bout frequency (15 to 18 s bin, p < 0.05; **Fig. 2G**; **Fig. S2E**). Separating bouts into turning and straight bouts showed that NEG, but not POS, stimuli increased turning bout frequency during the first 20 s after stimulus onset (NEG, p = 0.027; POS, p = 0.193; **Fig. 2H-J; Fig. S2F**), with only weak effects on straight bouts (**Fig. S2G,H**). Lower concentrations did not significantly alter turning bout frequency (**Fig. S2I**), consistent with the concentration dependence of posture angle responses.

Thus, partially immobilized larvae retained key valence-dependent behavioral features observed during free swimming. NEG stimuli increased swimming (bout frequency) and turning, whereas POS stimuli transiently reduced swimming. Because posture angle responses occurred across both stimulus classes, the opposite bout dynamics were unlikely to reflect differential stimulus detection or generalized motor activation alone (**Fig. 2D**; **Fig. S2E**). Thus, appetitive and aversive stimuli retained distinct motor signatures under imaging-compatible conditions.

### Chemosensory stimulation evoked widespread neuronal activity

To map chemosensory-evoked neuronal activity and relate it to motor output, we performed volumetric two-photon Ca²⁺ imaging in Tg(*elavl3:Hsa.H2B-GCaMP6s*) larvae ^52^, acquiring nine planes per volume at 3 Hz (**Fig. 3A**). We recorded sequential anterior and posterior fields of view (FOVs) in each fish, alternated acquisition order across fish, and registered all datasets to the mapzebrain reference brain ^53^. Across 10 fish, we extracted ∼200,000 ROIs, corresponding to ∼25% of neurons per fish on average ^54^.

**Figure 3.**
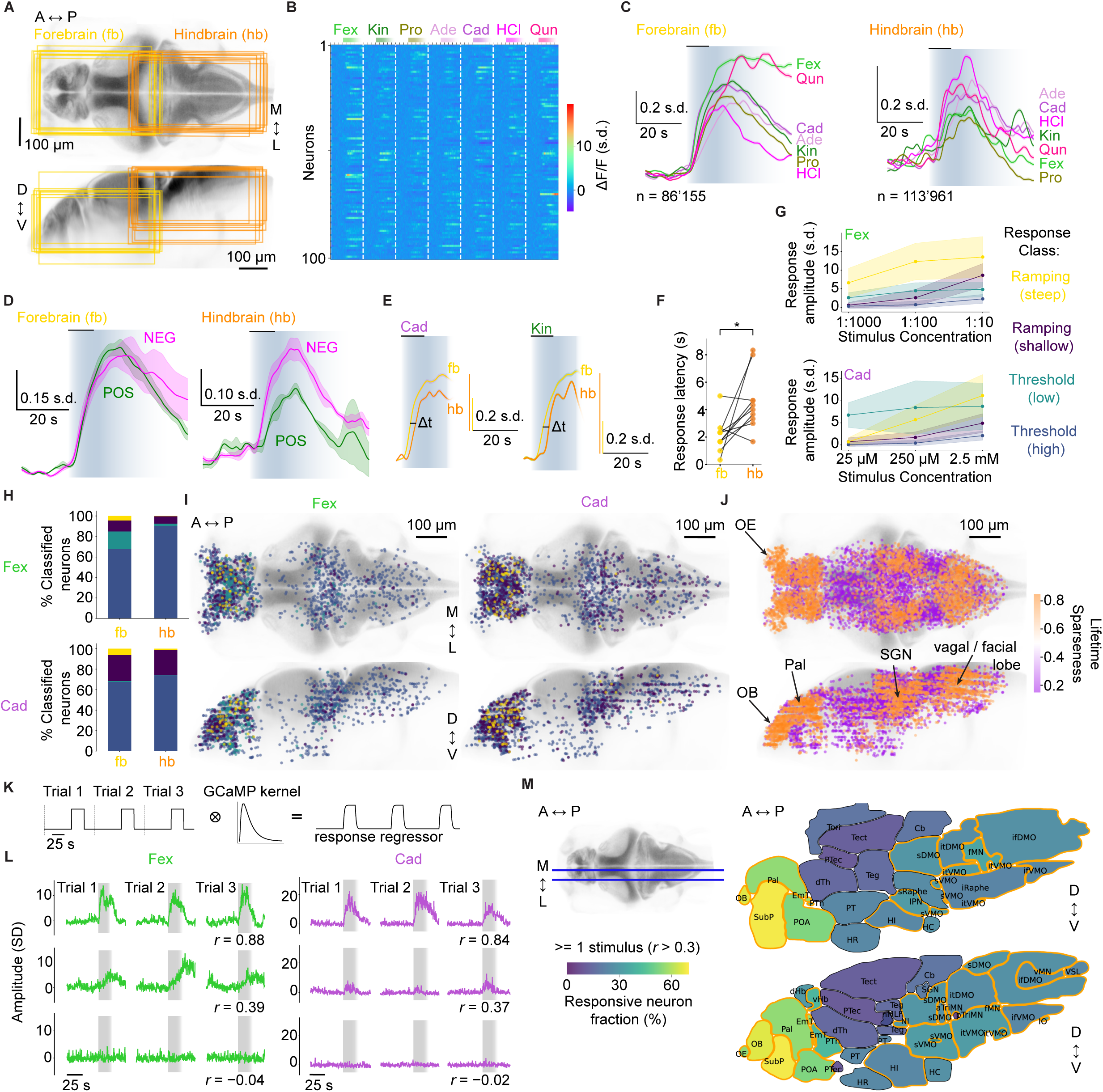
Chemosensory stimulation evokes widespread neuronal activity. (**A**) Dorsal (top) and lateral (bottom) views of the larval zebrafish brain showing field-of-view (FOV) outlines for all N = 10 fish (yellow, forebrain [fb] FOV; orange, hindbrain [hb] FOV). FOVs were aligned to a standard reference brain to illustrate spatial coverage across animals (**Methods**). (**B**) Raster plot of 100 randomly selected forebrain neurons showing trial-averaged F(Ca^2+^) traces (‘activity’) for the seven stimuli at their highest concentrations, z-scored relative to pre-stimulus baseline. (**C**) Trial-averaged z-scored neural activity traces from forebrain (left) and hindbrain (right) FOVs, showing responses to all seven stimuli (N = 10 fish). Background shading indicates approximate stimulus profile; trace shading indicates s.e.m. across neurons; traces are color-coded by stimulus identity. (**D**) Mean ± s.e.m. responses averaged across appetitive (POS; green) or aversive (NEG; magenta) stimuli in forebrain (left) and hindbrain (right) FOVs, illustrating region-specific, valence-dependent response differences. (**E**) Mean ± s.e.m. responses in forebrain and hindbrain FOVs for Cad and Kin, illustrating consistent response delays between FOVs. Y-axes are scaled separately to facilitate comparison of response dynamics. (**F**) Response latency in forebrain and hindbrain FOVs across all fish; Wilcoxon-Mann-Whitney test, p = 0.014. (**G**) Average response amplitudes of neurons grouped into four concentration-response clusters for Fex (green, top) and Cad (purple, bottom): cluster 1, steep-ramping (yellow); cluster 2, shallow-ramping (dark purple); cluster 3, low-threshold (turquoise); cluster 4, high-threshold (dark blue). (**H**) Proportions of each cluster type within forebrain and hindbrain FOVs for the two stimuli. (**I**) Anatomical distribution of classified neurons mapped onto the standard reference brain, color-coded by cluster identity; 1 in 10 neurons shown for clarity. (**J**) Spatial distribution of responsive neurons with the highest and lowest 2% lifetime sparseness values across the brain. Neurons are color-coded by lifetime sparseness (0-1); high sparseness (orange) indicates sharply tuned responses and low sparseness (purple) indicates broad responses. Olfactory epithelium (OE), olfactory bulb (OB), pallium (Pal), secondary gustatory nucleus (SGN), and vagal/facial lobe are indicated (arrows). (**K**) Binary stimulus regressors across three trials, with a 25-s response window used to capture slow stimulus washout. Regressors were convolved with a GCaMP kernel to model Ca²⁺ indicator kinetics (**Methods**). (**L**) Example neurons with high, moderate, or no correlation to the Fex (left) or Cad (right) stimulus regressors; Pearson’s *r* is shown below each trace. Grey shading shows the convolved stimulus regressor. Trials (stimulus repetitions) are shown concatenated, for visualization only, but were actually interleaved across blocks. (**M**) Left: reference medial and lateral slice positions for the anatomical maps. Right: fraction of neurons in each brain region with Pearson correlation coefficient *r* > 0.3 to any highest-concentration stimulus regressor, pooled across all 10 fish. Orange outlines denote regions with significantly more responsive neurons than expected by chance; binomial test, p < 0.05. A, anterior; P, posterior; M, medial; L, lateral; D, dorsal; V, ventral. All brain region abbreviations are defined in **Methods**.

Chemosensory stimulation elicited diverse response time courses across neuron-stimulus pairs (**Fig. 3B**). All stimuli produced clear responses of similar amplitude in the forebrain, whereas aversive stimuli elicited stronger average responses in the hindbrain (**Fig. 3C,D**). At the population level, hindbrain responses lagged behind forebrain responses by ∼2 s in sessions with counterbalanced acquisition order (p = 0.014; **Fig. 3E,F**). Response amplitudes depended on stimulus concentration (**Fig. 3G**). Clustering concentration-response profiles revealed a predominant high-threshold class, comprising more than 60% of responsive cells, that responded only at the highest tested concentration (**Fig. 3H,I**; **Fig. S3A-C**). Smaller ramping and low-threshold classes were also present, with ramping responses more common in the forebrain. Because behavioral responses were most robust at the highest concentrations, subsequent analyses focused on these conditions. Tuning sharpness, quantified by lifetime sparseness, was highest in canonical chemosensory regions, including the olfactory epithelium (OE), olfactory bulb (OB), and pallium (Pal), as well as gustation-related hindbrain nuclei such as the facial and vagal lobes and the secondary gustatory nucleus (SGN), while stimulus-responsive neurons were distributed throughout the brain (**Fig. 3J**).

We next quantified the fraction of responsive neurons in each annotated brain region using stimulus-specific regressors for the highest tested concentrations, classifying neurons as responsive when their correlation with the respective regressor exceeded *r* ≥ 0.3 (**Fig. 3K-M**; **Methods**). The highest responsive fractions occurred in forebrain regions with known olfactory function, including OE (55%), OB (66%), Pal (56%), and SubP (69%), and in several diencephalic regions, including preoptic area (POA; 53%), prethalamus (PTh; 42%), and eminentia thalami (EmT; 58%). Responsive neurons were detected in all regions but were less frequent elsewhere, particularly in the optic tectum (Tect; 12%) and pretectum (PTec; 11%). Across individual stimuli, the regional pattern was broadly conserved, although hindbrain responses differed more strongly between aversive and appetitive conditions (**Fig. S3D**). Similar regional distributions were obtained across correlation thresholds and when considering neurons responsive to more than one stimulus (**Fig. S3E,F**).

Taken together, chemosensory stimulation evoked widespread neuronal activity across the larval zebrafish brain. Responses were strongest and earliest in forebrain regions, whereas aversive cues elicited stronger and delayed responses in the hindbrain. Stimulus tuning was sharpest in canonical olfactory and gustatory regions, but responsive neurons were broadly distributed, with the highest fractions in telencephalic and diencephalic regions.

### Stimulus identity was decoded most accurately in olfactory bulb and pallium

Olfactory processing in zebrafish has been studied mainly in adults and in selected regions, especially OB, with more limited work in habenular, pallial, and subpallial areas ^55–59,31,60,61^. Because our recordings were performed in larvae and covered additional brain areas, we used stimulus identity coding as a benchmark for chemosensory processing across twelve highly responsive telencephalic and diencephalic regions. We quantified single-trial stimulus discriminability with a template-matching classifier (**Fig. 4A,B**; **Fig. S4A**; **Methods**), predicting stimulus identity from trial-wise population activity within each fish and region (**Fig. 4C**).

**Figure 4.**
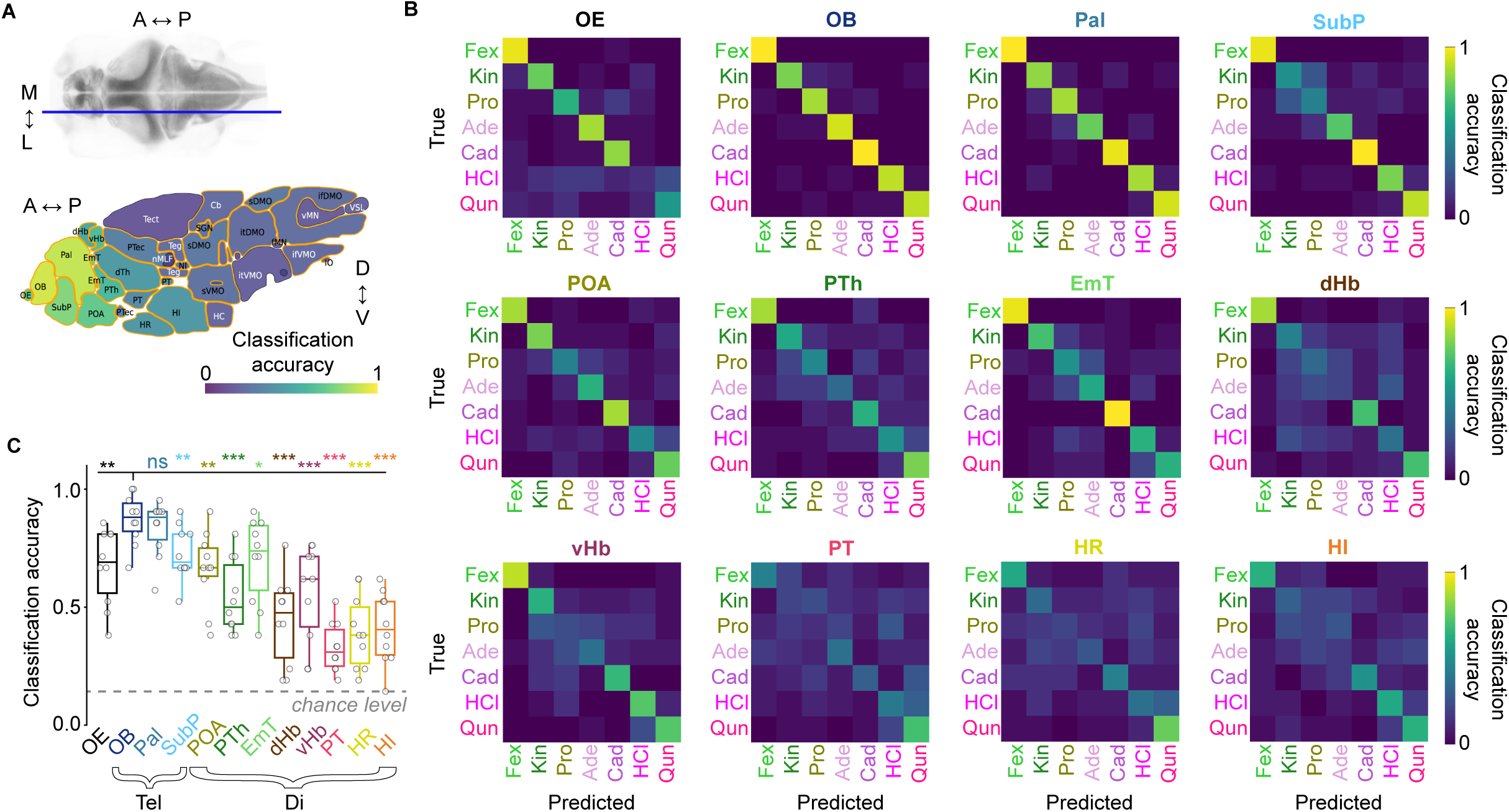
**Region-specific decoding of stimulus identity from population activity using template matching**. (**A**) Brain-wide map of stimulus-classification accuracy for each anatomical region, computed using a cross-validated template-matching algorithm and averaged across fish (N = 10). Orange outlines indicate regions with above-chance decoding performance; p-values were combined across fish using Fisher’s method, p < 0.05. (**B**) Confusion matrices for twelve selected telencephalic and diencephalic brain regions, averaged across fish. Rows indicate true stimulus identity and columns predicted identity; high diagonal entries indicate accurate classification. Color scale denotes normalized classification accuracy. (**C**) Classification accuracy across all regions shown in B. Asterisks denote significant differences relative to the olfactory bulb; Wilcoxon signed-rank test, *p < 0.05, **p < 0.01, ***p < 0.001; ns, not significant. Tel, telencephalon; Di, diencephalon. OE, olfactory epithelium; OB, olfactory bulb; Pal, pallium; SubP, subpallium; POA, preoptic area; Pth, prethalamus; EmT, eminentia thalami; dHb, dorsal habenula; vHb, ventral habenula; PT, posterior tuberculum; HR, rostral hypothalamus; HI, intermediate hypothalamus.

Decoding accuracy was highest in the OB (91 ± 7%, mean ± SD) and Pal (89 ± 9%), which contains the zebrafish homolog of olfactory cortex (Dp) ^62,48^ (**Fig. 4A-C**), but was significantly lower in the OE (68 ± 15%; Wilcoxon signed-rank test, p < 0.001). Several additional telencephalic and diencephalic regions showed intermediate accuracy, including SubP (76 ± 13%; p < 0.01), POA (69 ± 16%, p < 0.01), PTh (60 ± 13%, p < 0.001), EmT (72 ± 17%, p < 0.01), and the habenulae (vHb: 61 ± 14%, p < 0.001; dHb: 51 ± 16%, p < 0.001). In contrast, more posterior diencephalic regions, including the posterior tuberculum (PT; 36 ± 10%, p < 0.001), rostral hypothalamus (HR; 42 ± 17%, p < 0.001), and intermediate hypothalamus (HI; 43 ± 16%, p < 0.001), showed substantially lower classification performance despite clear stimulus-evoked responses (**Fig. S4B**). Classification accuracy was even lower throughout mid- and hindbrain areas (**Fig. 4A; Fig. S4C**).

To better understand regional differences in decoding accuracy, we quantified response similarity, trial-to-trial reliability, response latency, and tuning sharpness using lifetime sparseness (**Fig. S4D-G**). High-accuracy regions such as OB and Pal showed more separable (i.e. less similar) stimulus-evoked activity patterns, higher trial-to-trial reliability, shorter response latencies, and sharper tuning. By contrast, low-accuracy regions such as PT, HR, and HI showed more similar stimulus-evoked patterns, lower reliability, broader tuning, and delayed responses. On average, neurons in low-accuracy regions responded approximately 2 s later than OB neurons (**Fig. S4F**).

Overall, stimulus identity was reliably decodable in OB and Pal, moderately decodable in other telencephalic and anterior diencephalic regions, and weakly decodable in posterior diencephalic regions despite robust stimulus-evoked responses. This regional decline in decoding accuracy was associated with longer response latencies, lower trial-to-trial reliability, and broader tuning, consistent with less stimulus-specific chemosensory activity in posterior diencephalic areas. This organization was broadly consistent with observations from adult fish in previously studied regions ^63,56,57,59^.

### Small valence-related populations were regionally structured and functionally coupled

We next asked whether neuronal responses generalized across chemically distinct stimuli with similar behavioral effects. We defined valence-encoding neurons (VENs) as neurons whose activity aligned with appetitive (+VAL) or aversive (−VAL) stimulus groups rather than with individual chemical identities (**Fig. 5A**). HCl was excluded to balance the two groups, yielding three stimuli per class (NEG: Ade, Cad, Qun; POS: Fex, Kin, Pro; **Methods**). Neurons significantly correlated with either valence regressor after shuffle-based testing were classified as VENs (p_FDR_ < 0.05; **Fig. S5A**).

**Figure 5.**
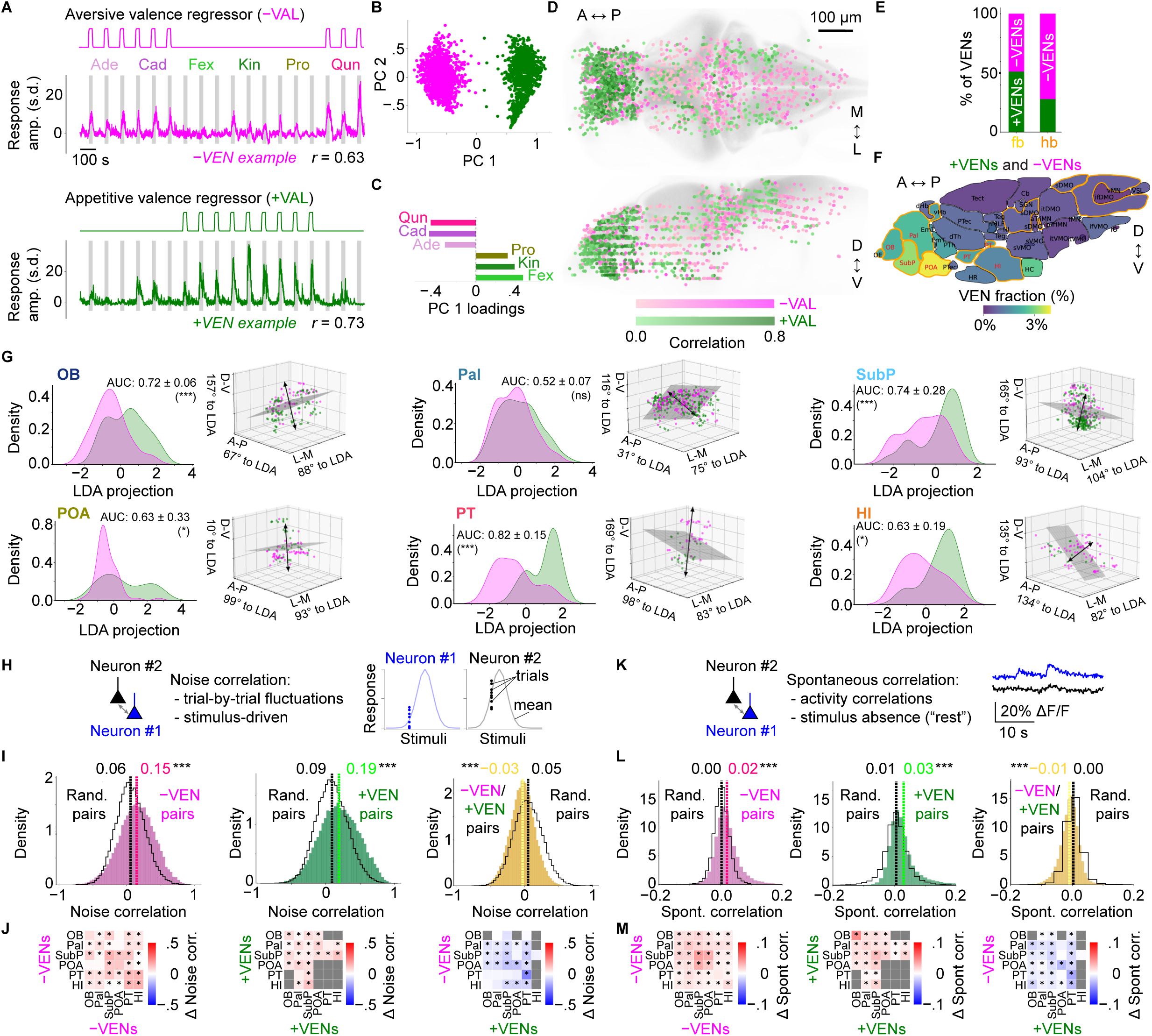
**Identification, spatial distribution, and functional coupling of valence-encoding neurons (VENs)**. (**A**) Aversive (−VAL, magenta) and appetitive (+VAL, green) valence regressors used to identify valence-encoding neurons (VENs), with example correlated neurons shown below. Trials are shown concatenated for visualization; actual stimulus repetitions were interleaved across blocks, with stimulus order randomized within each block. (**B**) Principal component (PC) projection of VEN response profiles; each dot represents one VEN. (**C**) PC 1 loadings showing that all stimuli contribute to PC 1, which separates valence. (**D**) Spatial distribution of −VENs and +VENs, color-coded by correlation with the corresponding valence regressor. (**E**) Proportions of −VENs and +VENs in forebrain and hindbrain FOVs. (**F**) Fraction of neurons significantly correlated with either valence regressor (p_FDR_ < 0.05), expressed as percentage of all neurons per brain region. Six forebrain regions with >50 VENs and at least 10 +VENs and 10 −VENs are highlighted in red. (**G**) LDA-based spatial separation of VENs across six brain regions. For each region, left panels show −VENs (magenta) and +VENs (green) projected onto the anatomical LDA axis, with larger area under the curve values (AUC; mean ± s.d.) indicating greater separation strength; permutation test with VEN label shuffling, *p < 0.05, ***p < 0.001; ns, not significant. Right panels show VEN positions in 3D, with the LDA vector (black arrow) and decision plane (grey). SubP, OB, POA, PT, and HI show pronounced dorsoventral or anterior-posterior separation, whereas Pal shows little separation. (**H**) Schematic of noise correlations, computed from trial-by-trial response fluctuations around mean stimulus responses. (**I**) Noise correlations distributions for −VEN/−VEN (magenta; n = 40,494), +VEN/+VEN (green; n = 61,843), and −VEN/+VEN pairs (yellow; n = 42,724), compared with area-and number-matched random pairs (black). Dashed lines indicate means; Wilcoxon-Mann-Whitney test, ***p < 0.001. (**J**) Mean noise correlation differences relative to matched random pairs within and across six forebrain valence regions; empirical p-values were combined across animals using Fisher’s method with FDR correction, *p < 0.05. Random-pair and unsubtracted VEN matrices are shown in **Fig. S5**. (**K**) Schematic of spontaneous correlations computed from pre-stimulus ΔF/F traces. (**L,M**) Same as I,J for spontaneous correlations; random-pair and unsubtracted VEN matrices are shown in **Fig. S5**. OB, olfactory bulb; Pal, pallium; SubP, subpallium; POA, preoptic area; PT, posterior tuberculum; HI, intermediate hypothalamus. Other abbreviations are defined in **Methods**. A, anterior; P, posterior; D, dorsal; V, ventral; M, medial; L, lateral.

We then confirmed that VEN classification reflected shared valence group rather than single-stimulus responses. In principal component space, −VENs and +VENs separated along the main axis (**Fig. 5B**), with comparable contributions from each stimulus to this separation (**Fig. 5C**). Clustering within each valence group further showed subclusters responding to multiple same-valence stimuli across fish (**Fig. S5B**). VENs were distributed across the brain but concentrated in forebrain regions (**Fig. 5D,E**). They represented only a small fraction of all neurons, peaking at ∼4% in POA and ∼3% in SubP, with both −VENs and +VENs showing clear regional bias (**Fig. 5F**; **Fig. S5C**).

In mammals, several olfactory areas, including the olfactory bulb and olfactory tubercle, show some spatial separation of domains associated with aversive and appetitive odor responses ^64,65^. We therefore asked whether +VENs and −VENs were spatially separated within VEN-rich regions. Across six regions containing at least 50 VENs (**Methods**), linear discriminant analysis (LDA) identified separating axes that often aligned with principal anatomical axes (**Fig. 5G**; **Fig. S5D**). Dorsoventral separation was evident in OB, SubP, POA, and PT, mediolateral separation in POA, and anterior-posterior separation in PT and HI, whereas Pal showed little clear topography, consistent with the proposed stochastic innervation of pallial neurons by olfactory bulb projection neurons in larval zebrafish ^62^. Thus, several VEN-rich regions showed spatial separation of +VENs and −VENs.

Because VENs were distributed across multiple regions, we next asked whether they showed coordinated activity across the six VEN-rich regions (**Fig. S5E**). Pairwise noise correlations, computed from trial-by-trial fluctuations in stimulus-evoked responses, and spontaneous correlations from pre-stimulus activity revealed a consistent sign structure (**Fig. 5H-M**): same-sign VEN pairs showed enriched positive noise correlations, whereas opposite-sign pairs showed enriched negative correlations relative to matched random neuron pairs (**Fig. 5I**). Region-resolved analyses showed broad −VEN coupling, with strongest enrichment between PT and HI and between SubP and POA, PT, and OB (**Fig. 5J**; **Fig. S5F**). +VEN coupling was more spatially restricted and particularly involved SubP and Pal. Opposite-sign negative correlations were strongest locally in PT, with additional cross-region negative correlations involving PT +VENs and −VENs in POA, SubP, and Pal. Spontaneous correlations broadly preserved this sign structure (**Fig. 5L**) but differed regionally: −VEN coupling was strongest in SubP and POA, whereas +VEN coupling was strongest in OB (**Fig. 5M**; **Fig. S5G**).

Same-sign coupling was stronger within than across regions, while opposite-sign correlations were similarly distributed within and across regions (**Fig. S5H**).

Overall, VENs formed small, forebrain-biased populations whose activity generalized across stimuli within behaviorally defined valence classes. These populations showed regional spatial organization, with separation of +VENs and −VENs in several regions, enriched positive correlations among same-sign VEN pairs, and enriched negative correlations among opposite-sign pairs.

### Behavior-correlated neurons were widespread, regionally enriched, and largely distinct from VENs

VENs captured activity aligned with behaviorally defined valence groups, but their overlap with motor-correlated neuronal populations remained unclear. We therefore used simultaneously recorded tail angle traces to define behavior-correlated neurons (BCNs) associated with distinct locomotor variables and compare their distribution with VENs.

Because chemosensory stimulation primarily modulated turning bouts in our assay (**Fig. 2**; **Fig. S2**), we first generated a turning bout regressor from asymmetric bouts convolved with a GCaMP kernel (**Fig. 6A**; **Methods**). Neurons significantly and strongly correlated with this regressor were classified as turn-BCNs (p_FDR_ < 0.05, *r* ≥ 0.3). Turn-BCNs were distributed across the brain but enriched in PT, nucleus of the medial longitudinal fasciculus (nMLF), tegmentum (Teg), nucleus isthmus (NI), and subdivisions of the medulla oblongata (MO; **Fig. 6B,C**), regions that generally contained few strongly correlated VENs (**Fig. S5C**). BCNs were also substantially more numerous than VENs: nMLF, which contains the anterior-most reticulospinal neurons ^50,66^, showed the highest turn-BCN fraction (∼29%), whereas strongly correlated VENs peaked at ∼2% in SubP (**Fig. 6D**; **Fig. S5C**).

**Figure 6.**
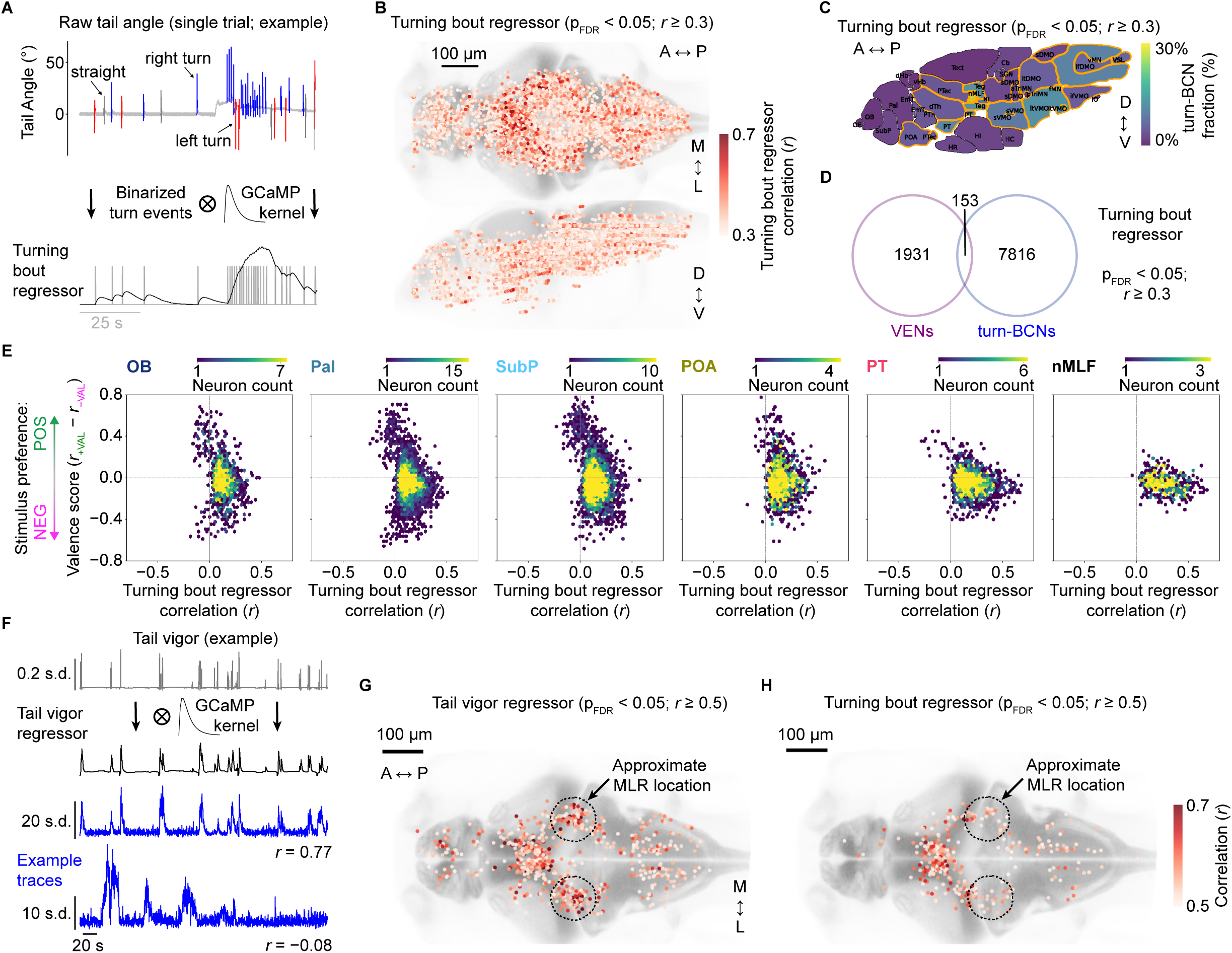
Identification of behaviorally correlated neurons (BCNs). (**A**) Turning bout regressor construction. Top: raw tail angle trace with swim bouts classified as straight (gray), leftward (red), or rightward (blue) turns. Bottom: turning bout times (gray) convolved with a GCaMP kernel to generate the turning bout regressor (black). (**B**) Spatial distribution of turn-BCNs, defined as neurons significantly and strongly positively correlated with the turning bout regressor (p_FDR_ < 0.05; *r* ≥ 0.3). Significance of individual neuron correlations was assessed by trial-shuffling (**Methods**). (**C**) Fraction of turn-BCNs across brain regions, pooled across fish; orange outlines indicate significant regional enrichment; binomial test, FDR-corrected p_FDR_ < 0.05. (**D**) Overlap between VENs (p_FDR_ < 0.05) and significant turn-BCNs (p_FDR_ < 0.05, *r* ≥ 0.3): 153 neurons, corresponding to 7% of VENs and 2% of turn-BCNs. (**E**) Joint distribution of valence score and turning bout regressor correlation across VENs and turn-BCNs in selected regions (VEN- or BCN-rich). Each plotted bin represents a combination of valence score (*r*(+VAL) − *r*(−VAL)) and Pearson correlation with the turning bout regressor, with color indicating the number of neurons in that bin. (**F**) Tail vigor regressor construction. Top: example tail vigor trace across trials. Middle: tail vigor regressor after convolution with a GCaMP kernel. Bottom: example neuronal traces with Pearson’s correlation coefficients. Scale bar, 20 s. (**G,H**) Spatial distribution of neurons highly correlated (*r* ≥ 0.5) with the tail vigor regressor (**G**) or turning bout regressor (**H**). Dotted circles indicate the approximate location of the mesencephalic locomotor region (MLR), a conserved premotor area involved in forward locomotion ^98^.

We next compared valence-related and turning-related activity at the single-neuron level. For each VEN and turn-BCN, we plotted its valence score, defined as *r*(+VAL) − *r*(−VAL), against its correlation with the turning bout regressor (**Fig. 6E**). In OB, Pal, and SubP, neurons spanned a broad range of valence scores but showed mostly weak turning correlations. In contrast, PT and especially nMLF neurons showed stronger turning correlations and a narrower spread along the valence axis. Thus, valence-related activity and turning bout correlation formed partly separable dimensions whose relative weighting changed across regions. In hindbrain regions, valence scores were less broadly distributed, biased toward negative values, and more tightly related to turning bout correlations (**Fig. S6A**). Thus, valence-related and turning-correlated activity were partly separable in forebrain regions but more closely aligned in hindbrain regions.

To test whether BCN maps were specific to turning or reflected broader motor output, we next analyzed correlations with a tail vigor regressor generated from tail angle variability convolved with a GCaMP kernel (**Fig. 6F**; Methods). Tail vigor revealed a broadly similar spatial pattern to the turning bout regressor (**Fig. S6B**), including enrichment in PT, but also highlighted a prepontine cluster near the midbrain-hindbrain boundary with particularly high correlations (*r* > 0.5; **Fig. 6G,H**). This cluster may correspond to the proposed zebrafish mesencephalic locomotor region (MLR) and largely overlapped with the mapzebrain NI annotation ^53,67^. As an additional comparison, a straight bout regressor identified fewer strongly correlated neurons, consistent with the weaker modulation of straight bouts in this assay (**Fig. S6C**; **Fig. S2**).

Thus, BCNs were widespread and enriched in motor-associated and premotor regions, whereas VENs formed smaller, anteriorly biased populations that were largely distinct from BCN populations. Continuous regressor comparisons showed that valence-related and movement-correlated activity were partly separable, especially in forebrain regions. PT contained both VENs and BCNs, identifying it as a candidate region for subsequent analyses of VEN-BCN coupling (**Fig. 5**; **Fig. 6**).

### Aversive and appetitive VENs show opposite coupling to BCNs

Having established that forebrain VENs and BCNs were largely distinct, and that valence-related and movement-aligned activity were partly separable, we next asked whether valence-behavior relationships differed between −VENs and +VENs. We therefore analyzed the two VEN classes separately, testing how their activity covaried with trial-by-trial locomotor expression, aligned with movement regressors, and correlated with BCNs.

We first asked whether VEN response amplitudes covaried with locomotor expression across trials. Trials were grouped as movement-increase, movement-decrease, or no-change trials (**Fig. 7A**; **Methods**). Pooling forebrain VENs, −VENs showed stronger responses in movement-increase trials and weaker responses in movement-decrease trials (**Fig. 7B,D**). In movement-decrease trials, −VEN activity was also reduced during appetitive stimulation, coinciding with the transient reduction in bout output observed for appetitive stimuli (**Fig. 2**). By contrast, +VENs showed more similar responses across movement-increase and movement-decrease trials (**Fig. 7C,E**). A behavioral (BHV) modulation index confirmed this sign-specific pattern: −VENs showed predominantly positive indices across regions, whereas +VENs were generally negative, with the strongest negative values in PT (**Fig. 7F,G**). Thus, −VEN and +VEN amplitudes covaried with trial-by-trial behavioral expression in opposite directions.

**Figure 7.**
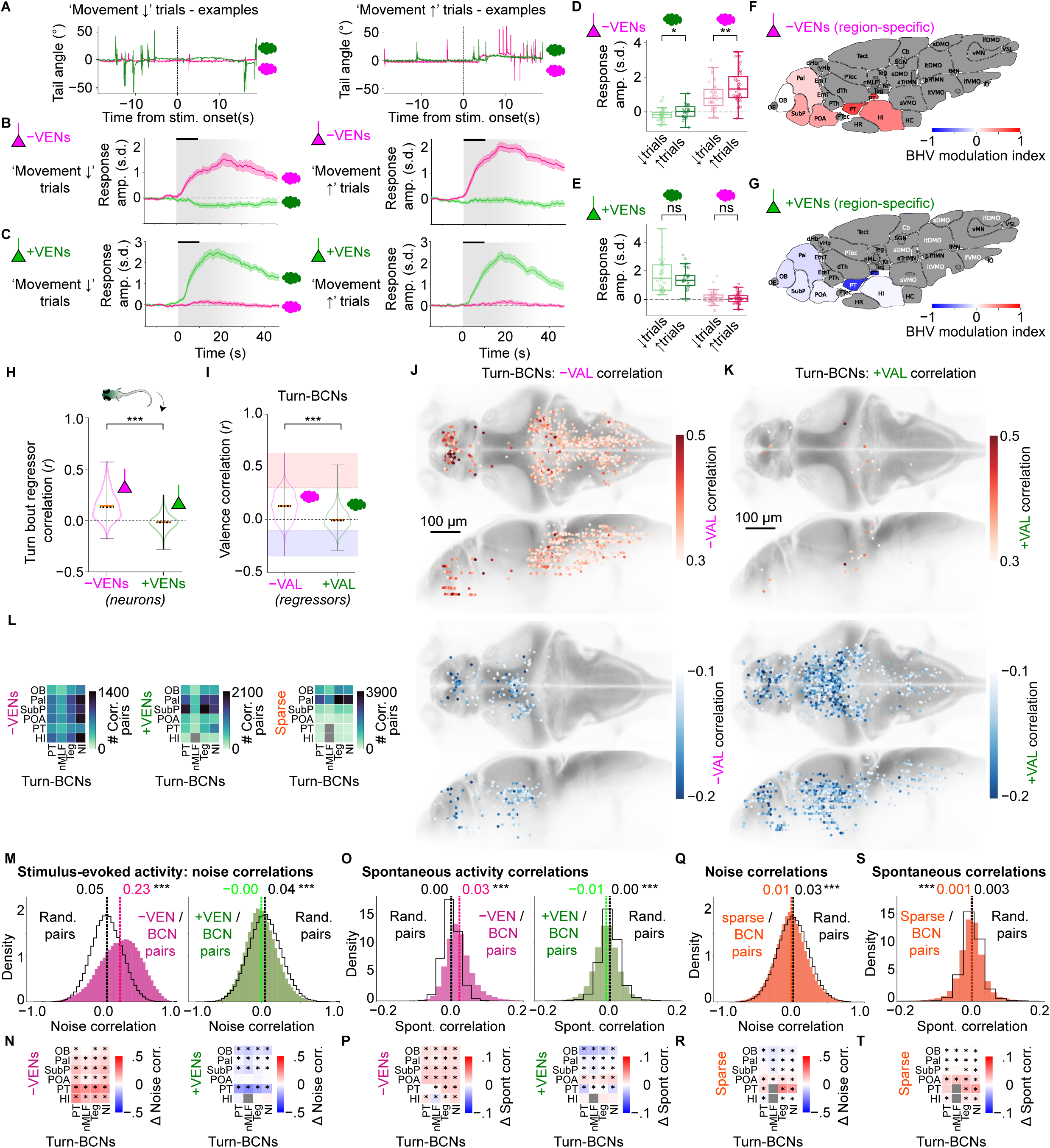
VENs show opposite correlations with behavioral activity. (**A**) Example raw tail angle traces for movement-decrease (‘Movement ↓’, left) trials and movement-increase (‘Movement ↑’, right) trials for an appetitive stimulus (Kin, green cloud) and an aversive stimulus (Qun, magenta cloud). (**B,C**) Mean ± s.e.m. of stimulus-evoked responses, averaged across z-scored neural activity traces, of forebrain −VENs (**B**) and +VENs (**C**), separated by movement-decrease and movement-increase trials, for appetitive (green cloud) and aversive (magenta cloud) stimuli, respectively. (**D,E**) Mean VEN activity per trial, quantified 15-25 s after stimulus onset and separated by stimulus valence and movement class, for −VENs (**D**; POS: p = 0.034, n↓ = 37, n↑ = 31; NEG: p = 0.0021, n↓ = 43, n↑ = 48) and +VENs (**E**; POS: p = 0.69; NEG: p = 0.46; same trial numbers; Wilcoxon-Mann-Whitney tests). (**F,G**) Behavioral (BHV) modulation index, ranging from −1 to +1, across six VEN-enriched regions for −VENs (**F**) and +VENs (**G**). (**H**) Correlations of −VENs and +VENs with the turning bout regressor; black dashed line, median; orange line, mean; ***p < 0.001. (**I**) Correlations of turn-BCNs with −VAL and +VAL regressors; ***p < 0.001. Shaded regions indicate the positively and negatively correlated turn-BCN subsets that are mapped anatomically in panels J and K. (**J,K**) Anatomical distribution of turn-BCNs with positive correlations (*r* ≥ 0.3; top, warm colors) or negative correlations (*r* ≤ −0.1; bottom, cool colors) to −VAL (**J**) or +VAL (**K**). (**L**) Average counts of VEN-turn-BCN and sparse neuron-turn-BCN pairs per fish for each region pair; gray squares mark region pairs with <100 average neuron pairs, which were excluded from further analysis. (**M**) Distributions of noise correlations for −VEN-turn-BCN pairs (magenta; n = 150,493) and +VEN-turn-BCN pairs (green; n = 167,451), compared with matched random pairs (black). (**N**) Region-specific differences in noise correlations between VEN-turn-BCN and matched random pairs for −VENs (left) and +VENs (right); significance was assessed by permutation testing with FDR correction. Gray squares indicate region pairs with <100 average pairs per fish. (**O,P**) Same as M,N for spontaneous correlations. (**Q,R**) Same as M,N for correlations between turn-BCNs and the top 2% most sharply tuned neurons (“sparse neurons”; n = 377,506 pairs). (**S,T**) Same as O,P for correlations between turn-BCNs and sparse neurons.

We next asked how the two VEN classes and turn-BCNs aligned with each other’s defining regressors. −VENs were generally positively correlated with the turning bout regressor, whereas +VENs showed weaker and slightly negative correlations (**Fig. 7H**). Conversely, turn-BCNs were positively correlated with −VAL but showed little positive correlation with +VAL (**Fig. 7I**). Strong −VAL-correlated turn-BCNs were concentrated near the midbrain-hindbrain boundary, in the hindbrain, and in SubP (**Fig. 7J**), whereas +VAL-correlated turn-BCNs were rare and many turn-BCNs were negatively correlated with +VAL (**Fig. 7K**). These +VAL-anti-correlated turn-BCNs were, as a notable contrast, mostly found in fore-and midbrain territories. Similar patterns were observed for the tail vigor regressor (**Fig. S7A-C**). Thus, regressor alignment was asymmetric, linking aversive valence more strongly to movement-correlated activity.

We then tested whether this alignment was reflected in pairwise VEN-BCN correlations (**Fig. 7L**). For stimulus-evoked responses, noise correlations with turn-BCNs diverged by valence sign: −VENs showed stronger correlations, whereas +VENs showed reduced correlations relative to matched random-pair controls (**Fig. 7M**). Similar patterns were observed for vigor-BCNs (**Fig. S7D**). Region-resolved analysis highlighted PT: −VEN-turn-BCN coupling was strongest for turn-BCNs in PT, albeit elevated across all six VEN-rich regions, whereas reduced +VEN-turn-BCN coupling was most prominent for PT +VENs (**Fig. 7N**). Spontaneous correlations preserved the same overall sign structure, but with a different regional pattern: −VEN coupling was strongest in OB, Pal, SubP, and POA, whereas reduced +VEN coupling was strongest in OB (**Fig. 7O,P**). Thus, sign-specific VEN-BCN relationships were partly present before stimulation, while PT was most prominent in stimulus-evoked response correlations.

Finally, we asked whether BCN coupling was specific to valence-related activity or also present in sharply stimulus-selective neurons. Applying the same pairwise correlation analysis to the 2% most sharply tuned neurons, defined by lifetime sparseness (**Fig. 3**), revealed no enhanced BCN coupling. Instead, sharply tuned neurons showed a small but significant reduction in BCN coupling for both noise and spontaneous correlations (p < 0.001; **Fig. 7Q-T**), even though it included neurons selective for both aversive and appetitive stimuli. Consistent with this, stimulus-specific analyses showed that forebrain neurons highly selective for the aversive stimuli Qun and Cad (*r* > 0.5), located mainly in OB, Pal, and SubP, also showed reduced coupling with BCNs (**Fig. S7F,G**). Thus, even aversive-selective stimulus-tuned neurons were not positively coupled to BCNs, indicating that BCN coupling was not a general property of stimulus responsiveness or identity selectivity.

Overall, these analyses revealed opposite relationships of −VENs and +VENs to behavioral output and BCN activity. −VENs were associated with increased locomotor expression, positive movement-regressor alignment, and enhanced BCN coupling, whereas +VENs were associated with reduced locomotor expression, weak or negative movement alignment, and reduced BCN coupling. This sign-specific pattern was not observed for sharply tuned stimulus-selective neurons, indicating that BCN coupling was more closely associated with valence-related activity than with stimulus identity coding.

## DISCUSSION

Our results show that behaviorally defined chemosensory valence is represented across a distributed brain-wide architecture in a naïve vertebrate brain. Rather than being confined to a single sensory region or motor-associated node, appetitive and aversive chemosensory stimuli engaged small, regionally structured valence-related populations across several forebrain regions. These populations were partly separable from movement-correlated neurons, especially in telencephalic regions, while posterior diencephalic and hindbrain regions showed tighter alignment with motor-correlated activity. Thus, larval zebrafish reveal a distributed brain-wide architecture in which chemosensory valence-related activity spans multiple circuit nodes and shows valence-specific relationships with motor output.

### Valence-dependent behavior is preserved across assay contexts

Across free-swimming and partially immobilized conditions, appetitive and aversive cues biased locomotion in qualitatively consistent ways. Aversive cues increased speed, turning, and bout frequency, whereas appetitive cues slowed freely swimming larvae near the stimulus source and transiently reduced bout frequency during imaging. Because posture angle responses, sustained orienting-like tail deflections, were comparable across both valence classes, differences in stimulus detection alone appeared insufficient to explain the divergent bout dynamics. The later increase in bout frequency during appetitive trials may instead reflect an offset-related or search-like state as stimulus levels decline.

These patterns closely match recent zebrafish studies showing that appetitive and aversive chemical cues bias behavior toward distinct motor strategies. In free-swimming assays, aversive cues promote dispersal, turning, or escape-like locomotion, whereas attractive odors are associated with slower, more localized sampling ^13,34^, closely paralleling the opposing speed and turning effects observed here. Similarly, our partially immobilized data, in which aversive stimuli preferentially increased turning bouts, align with recent work linking avoidance-associated cues to turn-like motor states ^34^.

Similar principles are seen across insects and rodents, where attractive odors often slow movement and promote local search, whereas aversive odors increase avoidance-related locomotion ^68,69^. Thus, our behavioral results fit a broader framework in which appetitive and aversive cues bias distinct components of chemosensory navigation, including local sampling, forward progression, and avoidance-related reorientation ^5,70,34^.

### Brain-wide responses reveal region-specific stimulus representations

Chemosensory stimulation evoked widespread but regionally structured brain activity. Canonical olfactory regions, especially olfactory bulb and pallium, the dorsal telencephalic territory that includes the fish homolog of olfactory cortex (Dp) ^48,71,72^, carried the most separable and reliable stimulus identity representations. This is consistent with adult zebrafish studies identifying the olfactory bulb and the pallial subdivision Dp as prominent sites of odor identity coding ^55–57^, and suggests that key regional features of olfactory functional organization are already present early in development ^73,62^. By contrast, several posterior diencephalic (incl. hypothalamic) regions responded robustly but carried weaker identity information, consistent with more integrated response formats, possibly combining chemosensory, state-, or motor-related variables ^74,75^. Thus, larval chemosensory processing recruited a brain-wide network beyond early olfactory centers, while exhibiting regional features broadly aligned with adult zebrafish olfactory organization ^76,62^.

### Valence-related activity is distributed across small, coordinated neuronal populations

Using behaviorally defined stimulus classes, we identified small VEN populations whose activity generalized across appetitive or aversive cues rather than individual stimulus identities. A central finding was that this valence-related activity was partly separable from movement-correlated activity, even though valence classes were defined by behavior and aversive stimuli increased locomotor output. VENs and BCNs also overlapped only partly, and their regional distributions and coupling patterns differed, indicating that VENs were not simply movement-correlated neurons relabeled by stimulus class. Thus, VENs should be understood as operationally defined populations that capture stimulus-class-related activity partly separable from, but still interacting with, behavioral output. These populations were distributed across multiple brain regions, showing that valence-related activity was not confined to a single focal node even in a naïve vertebrate brain early in development. Here, we use “naïve” operationally to refer to larvae tested without explicit conditioning or stimulus-reward training.

VENs also showed local spatial organization. In several forebrain regions, +VENs and −VENs occupied partially distinct spatial domains, whereas pallial VENs were more dispersed, consistent with distributed olfactory projections to the pallium and with dispersed odor-specific ensembles in adult Dp ^57,62^. These spatial patterns could reflect finer substructure within larger annotated regions, such as amygdalar-, striatal-, or septopallidal-like subdivisions within the subpallium ^77,48,78^, supporting fine-grained organization of valence-related activity without necessarily implying a canonical valence map.

Beyond their anatomical organization, VENs showed structured functional relationships. Same-sign VEN pairs showed enriched positive correlations relative to matched random neuron pairs, whereas opposite-sign pairs showed enriched negative correlations, both during stimulation and before stimulus onset. This suggests that valence-related populations were embedded in a functional architecture that is not only being imposed by sensory drive. The opposite-sign relationships are broadly reminiscent of opponent motifs in Drosophila valence circuits ^79,80^, but testing whether they may reflect direct synaptic opponency will require approaches beyond correlation analyses.

Combined with the broader spatial distribution of −VENs compared with +VENs, the correlation analyses suggest that aversive cues engaged a more extended valence-related network, at least within the present stimulus set. More broadly, these findings, together with accompanying novel olfactory bulb work ^13^, support a multistage view in which preference-biased representations can arise in early olfactory circuits and become embedded within broader telencephalic and diencephalic networks. Such distributed networks could provide substrates for state- or experience-dependent modulation, as, for instance, suggested by learning-dependent reorganization of valence-related representations in the adult olfactory pallium ^31^.

This distributed organization aligns with the broader vertebrate view in which olfactory valence is associated with interacting sensory, evaluative, motivational, and action-related circuits rather than a single valence center. In mammals, innate and learned odor value have been linked to distributed olfactory bulb ^10–13^, amygdalar ^16–20^, olfactory tubercle/striatal ^21–25^, hypothalamic ^26–29^, and prefrontal/orbitofrontal networks ^17,14,30^. In larval zebrafish, brain-wide responses to alarm cues and cadaverine have likewise shown that aversive chemosensory stimulation recruits distributed networks beyond early olfactory centers ^32,33^. Building on these studies, our dataset maps responses to a chemically diverse, behaviorally validated panel of appetitive and aversive chemosensory cues into a common brain-wide framework, combining cellular-resolution activity imaging with simultaneous motor readout in the same preparation. This allowed us to relate distributed valence-related activity directly to stimulus identity representations and movement-correlated populations.

### Valence-related and movement-correlated signals are sign-specifically coupled

Although VENs and BCNs were partly separable, their relationship varied by region and valence sign. BCN coupling was generally more closely associated with valence-related activity than with sharply tuned stimulus identity coding. In OB, Pal, and SubP, individual neurons were broadly distributed along the valence axis but showed weak turning correlations, indicating greater separability from movement. By contrast, PT, nMLF, and hindbrain regions showed stronger turning correlations and a more restricted neuronal distribution along the valence axis, consistent with tighter valence-movement alignment in posterior motor-associated circuits. This suggests that anterior VEN populations may be more closely associated with stimulus classification or evaluative components, whereas posterior populations may be closer to movement-related state changes and premotor circuits, including nMLF and hindbrain pathways ^50,74,75^.

Within this organization, +VENs were mainly associated with reduced or negatively coupled movement-correlated activity in diencephalic and midbrain-associated regions, whereas aversive stimuli more strongly recruited posterior motor-associated BCNs, including hindbrain regions, where stimulus identity decoding was weak and valence-related activity was more closely linked to turning behavior. Thus, appetitive and aversive valence-related activity did not appear to converge onto a single common motor-linked axis; instead, +VENs and −VENs were associated with sign-specific network relationships linked respectively to reduced versus enhanced movement-correlated activity. Because these relationships were derived from partially immobilized larvae, they capture only a limited part of the sensorimotor architecture. Nevertheless, the congruence between free-swimming and partially immobilized behavioral signatures suggests that these relationships reflect behaviorally relevant components of chemosensory navigation.

### Posterior tuberculum as candidate interface between valence-related and motor-linked signals

Among the regions identified here, the posterior tuberculum emerged as a prominent candidate interface between valence-related and motor-correlated activity. PT receives olfactory bulb projections in zebrafish ^62^ and, in our dataset, combined weak stimulus-identity decoding with valence-related spatial organization, BCN enrichment, and strong sign-specific VEN-BCN relationships. This profile suggests that PT may occupy a position within the distributed valence-related network where stimulus-class signals are more closely associated with motor-linked activity than in identity-rich olfactory regions.

This interpretation is consistent with comparative evidence that PT-like diencephalic systems contribute to odor-motor coupling and locomotor control in other vertebrates ^81–83^. The presence of dopaminergic neurons in PT ^47^ is also notable, given the broad involvement of dopamine in valence processing, locomotor control, and behavioral state regulation ^84,7,85^. However, our imaging experiments did not identify or manipulate dopaminergic PT neurons directly, and the cellular identity of PT VENs remains unresolved at the moment. Future studies will be needed to test whether PT contributes causally to valence-dependent locomotor changes and which PT cell types mediate this candidate valence-motor convergence.

Other regions may serve partially different functions within the same distributed architecture. Subpallium and preoptic area were enriched in VENs and contributed prominently to VEN coupling, consistent with roles in valence-related processing and state-dependent modulation ^74,86^. By contrast, although habenular circuits have been implicated prominently in odor-guided behavior, sensory-history integration, and action selection ^41,46,70^, the habenula did not emerge here as a dominant node in acute valence-motor coupling. This may indicate that habenular circuits are more strongly recruited during sensory-history-dependent, learned, state-dependent, or longer-timescale behavioral regulation than during the acute stimulus-movement relationships emphasized here ^87,88^. Together, these observations suggest that the acute valence-movement relationships captured here represent one layer of a broader, context-dependent architecture for chemosensory navigation.

## Conclusions

Together, these findings show that behaviorally defined chemosensory valence is represented across a distributed brain-wide architecture in a naïve vertebrate. Rather than being localized to a single sensory, evaluative, or motor-associated region, valence-related activity was carried by small, regionally structured populations that were partly separable from both stimulus identity coding and movement-correlated activity. Thus, even early-life chemosensory behaviors appear to involve distributed interactions among sensory, telencephalic, diencephalic, and motor-associated circuits.

Future work should test the causal roles of these candidate nodes and determine how internal state, experience, stimulus intensity, and active navigation shape this organization during flexible chemosensory-guided behavior.

## ACKNOWLEDGMENTS

We thank Krasimir Slanchev and Karin-Finger-Baier for excellent assistance with fish husbandry. We thank Thomas Offner for help with stimulus calibrations. We thank Oded Mayseless, Claire Meissner-Bernard, Thomas Offner, and Carlos Gabriel Aguilar Perez for critical comments on the manuscript and members of the Frank group for insightful discussions. We also thank Herwig Baier, Ilona Grunwald-Kadow, and Laura Busse for advice throughout this project. This work was funded by the Max Planck Society and the German Research Foundation (DFG) under Germany’s Excellence Strategy (EXC 2067/1390729940).

## AUTHOR CONTRIBUTIONS

Conceptualization, T.F. and B.J.; experimental design, T.F. and B.J.; experiments, B.J.; data analysis, B.J.; writing, review and editing, T.F. and B.J.; funding acquisition, T.F. supervision, T.F.

## DECLARATION OF GENERATIVE AI AND AI-ASSISTED TECHNOLOGIES IN THE WRITING PROCESS

During the preparation of this work, the authors used ChatGPT (OpenAI) to assist with language editing and code development. After using these tools, the authors carefully reviewed and edited all outputs and take full responsibility for the content of the published article. All scientific concepts, analyses, interpretations, and conclusions were conceived, implemented, and verified by the authors.

## METHODS

### Zebrafish husbandry

Larval zebrafish (*Danio rerio*) were raised in facility water according to established husbandry protocols. All animal procedures conformed to institutional guidelines of the Max Planck Society and were approved under licenses from the regional government of Upper Bavaria (ROB-55.2-2532.Vet_02-21-104, ROB-55.2-2532.Vet_02-22-29). Adult zebrafish were housed in a recirculating aquatic facility maintained at pH 7.0-7.5 and conductivity of 750-800 µS. Adult fish were fed Gemma Micron 300 using automated feeders two to three times per day and *Artemia salina* once daily. To obtain larvae for experiments, three males and three females were placed in sloped breeding tanks overnight. Eggs were collected the next morning, rinsed, and transferred to 92 × 16 mm Petri dishes in incubators maintained at 29 ± 1°C under a 14 h/10 h light/dark cycle. Imaging and behavior experiments were performed in larvae of undetermined sex, mostly at 7 dpf (days post fertilisation; age range in freely swimming experiments was 4-7 dpf). Larvae were fed Sera Micron Nature from 5 dpf onward, and Petri dish water was exchanged daily after feeding commenced.

For free-swimming behavioral experiments, Tg(*dlx5a-dlx6a:ITETA*)_fmi8Tg_; Tg(*5xUAS:iteta, ptet:gcamp5G, myl7:EGFP*) larvae were used because of their pigmentation and reliable breeding. For functional imaging experiments, Tg(*elavl3:Hsa.H2B-GCaMP6s*) larvae were used ^52^. At 3 dpf, embryos were lightly sedated with MS-222 and screened for fluorescence using an epifluorescence microscope. GCaMP6s-positive animals were selected for experiments.

### Behavioral experiments

#### Free-swimming behavioral experiments

Free-swimming preference assays were performed in custom-built rectangular acrylic arenas measuring 16 cm × 6.5 cm × 3.5 cm. Arenas contained slits at both ends for removable barriers that separated the larval swimming area from agarose stimulus patches during habituation.

Chemosensory stimuli were delivered from 3% agarose (Sigma-Aldrich, A9539) patches prepared in artificial fish water (AFW). AFW was prepared fresh weekly using 4 g ocean salt, 1.05 g NaHCO₃, and 0.12 g NaH₂PO₄ in 10 L deionized water, adjusted to pH 7.5.

For each experiment, the control side of the arena contained a 3% agarose patch prepared in AFW. On the stimulus side, the respective chemical stimulus was added to molten 3% agarose at 40-50°C before the patch was poured. Mixing at this temperature allowed the stimulus to dissolve into the agarose while avoiding thermal degradation. For control experiments, empty 3% agarose patches were placed on both sides of the arena. Experiments were performed at room temperature.

Arenas were placed inside a closed box to visually shield larvae from the surrounding environment and were illuminated from below using a diffuse LED white light source. Each arena was filled with 100 ml AFW, and typically 20 larvae were transferred individually using a pipette. Behavior was recorded from above at 30 Hz using a webcam (Logitech c920). After a 5 min habituation period, both agarose-patch barriers were removed simultaneously, and recording continued for at least 10 min.

Each free-swimming experiment was defined as one arena recording of a larval cohort under one agarose-patch condition. In stimulus experiments, one side of the arena contained a stimulus-loaded agarose patch and the opposite side contained an AFW-only control patch; in control experiments, both sides contained AFW-only agarose patches. Typically, approximately 20 larvae were tested per experiment, and each stimulus was assessed across 8-12 independent arena recordings. To control for day-to-day variation, an AFW-only control experiment was performed on each experimental day using larvae from the same cohort. Within a given experimental day, larvae could be exposed to up to three chemosensory stimulus conditions and the daily control condition, with stimulus and control order randomized across the day. Consequently, some larvae experienced both appetitive and aversive stimuli, whereas others experienced stimuli from only one behavioral class. Larvae were not reused across days. Thus, each experiment corresponded to a single stimulus condition in one arena recording. Some individual larvae could contribute to more than one stimulus condition within the same day; analyses were therefore performed at the stimulus-condition level, pooling larvae within each condition, and at the experiment level, using independent arena recordings.

### Chemosensory stimuli for the free-swimming assay

Stimulus diffusion in the arena was estimated using methylene blue (**Fig. S1A-D**). During the habituation period, no dye leakage from the agarose patch into the arena was observed, confirming that larvae were not exposed to the stimulus before barrier removal. After barrier removal, dye initially entered the arena through a variable convection phase, likely caused by turbulence during barrier removal. This was followed by slower diffusion across the arena. After 20 min, dye had not reached the far side of the arena, preserving a dye-free control region. However, we could not obtain per-stimulus, time-resolved concentration fields; therefore, compound-specific plume differences may contribute to variability observed in free-swimming assays.

To estimate the dye concentration near the source, average pixel intensity in the source-proximal region was compared with calibration solutions of known methylene blue concentration. This analysis indicated that the source-proximal concentration was diluted by approximately 500-fold after a 10 min gradient development period (**Fig. S1D**). Accordingly, high initial stimulus concentrations were used in the agarose patches. Because chemosensory compounds differ from methylene blue in molecular weight, this analysis was used only as an approximate estimate of stimulus distribution.

Stimuli used in the free-swimming assay were prepared as follows:

| Stimulus | Preparation | Final agarose-patch concentration |
| --- | --- | --- |
| Food extract (Fex) | 0.4 g Sera Micron powder in 100 ml | 1:10 dilution |
| Kin water (Kin) | 2 embryos/ml in AFW overnight (after <sup>86</sup>) | 1:1 dilution |
| L-proline (Pro) | 1 M stock in dH<sub>2</sub>O (Sigma-Aldrich, 81709) | 100 mM |
| Cadaverine dihydrochloride (Cad) | 200 mM stock in dH<sub>2</sub>O (Sigma-Aldrich, C8561) | 25 mM |
| Hydrochloric acid (HCl) | 1 M stock in dH<sub>2</sub>O (Sigma-Aldrich, H1758) | 56 mM |
| Quinine hydrochloride dihydrate (Qun) | 1.985g in 50 ml 3% agarose (Sigm-Aldrich, Q1125) | 100 mM |
| Adenosine | 25 mM stock in dH<sub>2</sub>O (Sigma-Aldrich, A9251) | 2.5 mM |

Our assay did not attempt to isolate olfactory from gustatory or trigeminal pathways. Because this screen was designed to identify stimuli that evoke robust population-level attraction or avoidance rather than to determine full concentration-response functions, classification as aversive (NEG) or appetitive (POS) refer to the behavioral effects observed under the conditions tested here.

### Behavioral tracking

Videos were cropped to the arena boundaries and trimmed to the 10-min period following barrier removal. Individual larvae were tracked using TRex ^89^. Although identity tracking was generally stable, occasional identity swaps occurred when larvae overlapped or moved in close proximity. To minimize tracking artifacts, only larvae tracked successfully for at least 90% of the experiment were included. In addition, speed values exceeding 30 mm/s were excluded to remove artifacts caused by rare identity swaps across the arena. To prevent static or minimally active larvae from biasing movement and behavioral analyses, the distribution of total distance travelled was inspected, and larvae moving less than 60 cm over the 10-min recording were excluded.

### Free-swimming behavioral measures

Behavioral metrics were extracted from tracking data to quantify responses to chemosensory stimuli. Analyses were restricted to the final 5 min after barrier removal, during which the stimulus gradient was more developed. Stimulus preference was calculated within the stimulus zone, defined as the half of the arena closest to the stimulus source. Other metrics (Δ heading angle, swim speed) were calculated within a narrower stimulus zone, defined as the 25% of the arena closest to the stimulus source.

Preference was quantified from the spatial distribution of larvae across the arena. The preference index was calculated from the relative occupancy of the stimulus and control zones, providing a measure of attraction or avoidance. Positive values indicated preference for the stimulus side, whereas negative values indicated avoidance. Additional metrics included mean x-position, swim speed, turning behavior, and spatial distribution within the arena. These metrics were used to compare stimulus-evoked behavioral changes across appetitive and aversive conditions. Differential occupancy (Δ occupancy) maps shown in **Fig. 1**, were calculated as follows: for each spatial bin, we first calculated the fraction of total larval occupancy in that bin. We then subtracted the corresponding normalized occupancy from the control experiments from the normalized occupancy in stimulus experiments. Positive values indicate greater occupancy than in control experiments, negative values indicate lower occupancy.

### Partially immobilized preparation

For partially immobilized recordings, awake, unanesthetized larvae were embedded dorsal side up in 2% low-melting-point agarose (Sigma-Aldrich, A9414) in the lid of a 35 mm Petri dish. After the agarose solidified, the dish lid was filled with AFW. The head and trunk were held in agarose, while agarose around the mouth/nasal region was removed to allow exposure to chemosensory stimuli.

The tail was also freed from the agarose to permit stimulus-evoked movements during simultaneous behavioral tracking and two-photon imaging. Tail movements were recorded at 60.85 Hz using a digital camera (Thorlabs) equipped with an IR filter and positioned below the fish, with illumination provided by an IR LED (870 nm; Thorlabs).

### Tail tracking

Tail movements were tracked using a custom tool based on the method of ^90^. Each video frame was divided by a background image to enhance contrast, and contours were extracted by thresholding using OpenCV. The algorithm identified the fish outline and internal landmarks, including the swim bladder. The fish heading was defined as the vector from the swim bladder to the midpoint between the eyes. Tail tracking began at the swim bladder and followed the longest skeletonized path to the tail tip. Evenly spaced points were interpolated along this path. The tail-tip angle was defined as the angle between the fish heading vector and the vector from the swim bladder to the tail tip.

### Identification of swim bouts and postural reorientation

Raw tail-angle traces were first zero-centered to correct for offsets introduced during embedding. Swim bouts were detected by calculating tail vigor as the standard deviation of the tail-angle trace in a rolling 100 ms window. Bouts were identified using a threshold of 0.1. A minimum bout duration was imposed to avoid false detections, and brief dips below threshold within a bout were ignored to prevent sustained movements from being split into multiple events. Bout start and end points were expanded by two frames to capture the full extent of each movement. Postural reorientation was quantified from the mean tail angle during periods before and after stimulus onset. This allowed slow stimulus-evoked deviations in tail posture to be analyzed separately from discrete swim bouts.

### Identification of lateralized bouts

Swim bouts were classified as left-turning, right-turning, or straight using a laterality index. For each bout, the tail-angle trace was extracted from bout onset for up to 70 ms, corresponding to four to five frames at 60.85 Hz, or for the full bout duration if the bout was shorter. This time window was selected because early bout kinematics correlate with the direction ultimately turned towards by freely swimming fish ^91,92^. The circular mean of the tail angle during this initial bout period was calculated for each bout. Bout laterality was then classified using a Gaussian mixture model fitted to the distribution of laterality indices. This approach allowed bouts to be separated into left-biased, right-biased, and non-lateralized categories.

### Bout and postural analysis

Stimulus-evoked changes in swimming activity were quantified from peristimulus time histograms of bout rate. Bouts were aligned to stimulus onset, and bout frequency was calculated in 3-s bins across a 20-s pre-stimulus and 30-s post-stimulus window. To account for inter-animal differences in baseline activity, bout frequencies were z-scored relative to each fish’s own pre-stimulus baseline.

Based on these PSTHs, bout metrics were quantified within 20-s pre- and post-stimulus windows relative to stimulus onset. Stimulus-evoked changes were tested using linear mixed-effects models with stimulus condition, defined as pre- versus post-stimulus, as a fixed effect, and fish identity and imaging session, corresponding to forebrain or hindbrain FOV, as random effects. For example, turning bout frequency was modeled as:

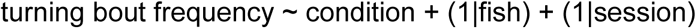

Statistical significance was determined from the fixed effect of stimulus condition. Linear mixed-effects models were implemented in Python using the statsmodels package.

Postural traces were derived from tail-angle recordings. Raw traces were first zero-centered to correct for embedding-related offsets. Swim bouts were identified from tail vigor, defined as the standard deviation of the tail angle in a rolling 100-ms window. To isolate slow postural reorientation from discrete swimming movements, detected bout periods, extended by 10 frames after bout offset to include residual motion, were linearly interpolated between the pre- and post-bout tail angles. The resulting continuous trace was low-pass filtered with a second-order Butterworth filter with a 0.5-Hz cutoff and baseline-corrected by subtracting the mean pre-stimulus tail angle. Postural responses were aligned to stimulus onset and quantified as the absolute deviation from baseline, thereby measuring sustained tail deflection independently of direction. Mean posture was compared between 20-s pre- and post-stimulus windows for each stimulus condition.

### Stimulus application protocol

For head-fixed behavioral and functional imaging experiments, chemosensory stimuli were delivered using a perfusion system consisting of two computer-controlled four-channel peristaltic pumps (Ismatec Reglo). The perfusion outlet was positioned near the larva’s mouth and nose, providing continuous AFW flow. During stimulation, flow was partially redirected to a stimulus channel while maintaining the same total flow rate, preventing mechanosensory responses caused by changes in flow. Tubing length was kept minimal and identical across all eight stimulus channels, which were combined into a common perfusion line using an 8-to-1 manifold before reaching the larva.

Computer-controlled switching of individual pump channels, synchronized to the imaging system, ensured that all stimuli shared the same final delivery path and avoided channel-specific differences in stimulus arrival time. Excess solution was removed through a vacuum line inserted into a small opening made in the agarose behind the larva.

Stimuli were presented in repeated trials. Each trial contained a baseline period (40 s), a stimulus period (10 s), and a washout period (50 s). Experiments using a fluorescent dye (Fluorescein; Sigma-Adrich) revealed an approximate 13-second delay between channel switching and stimulus arrival at the larva’s nose, such that the stimulus effectively reached the animal at 53 seconds from trial start. The solution reaching the larva was at room temperature.

Each larva underwent two imaging sessions, covering anterior and posterior field of view (FOV) in counterbalanced order across fish (see below). In each session, larvae received the same 15 stimulus-concentration conditions: all seven stimuli at their highest concentration, with Fex, Pro, Cad, and Qun additionally presented at two lower concentrations (100%, 10%, and 1% of the respective reservoir concentration; see table below). These conditions were delivered in three interleaved blocks per field of view, with each condition appearing once per block and the order randomized within each block, minimizing potential confounds from trial order. Thus, in each fish, each stimulus-concentration condition was presented three times per FOV and six times per fish in total. Stimuli and concentrations used for head-fixed behavioral and functional imaging experiments were:

| Stimulus | Reservoir concentration | Final concentration applied |
| --- | --- | --- |
| Food extract (Fex) | 1:10 | 1:10, 1:100, 1:1000 |
| Kin water (Kin) | 1:100 | 1:100 |
| Proline (Pro) | 2.5 mM | 2.5 mM, 250 μM, 25 μM |
| Adenosine (Ade) | 25 μM | 25 μM |
| Cadaverine (Cad) | 2.5 mM | 2.5 mM, 250 μM, 25 μM |
| HCl (pH5) | 1.27 mM | measured pH (in AFW) ∼ 5 |
| Quinine (Qun) | 2.5 mM | 2.5 mM, 250 μM, 25 μM |

This stimulus panel was selected to elicit robust appetitive or aversive behavior rather than to systematically sample intensity- or salience-matched chemosensory space.

### Two-photon Ca^2+^ imaging Image acquisition

Ca^2+^ imaging was performed using a two-photon movable objective microscope (MOM; Sutter Instruments) equipped with a 20x water-immersion objective (NA 1.0, Zeiss). Image acquisition and microscope control were performed using ScanImage software (Vidrio Technologies). Two-photon excitation was provided at 960 nm using a Ti:sapphire laser (Mai Tai; Spectra Physics). Emission was collected through a GaAsP photomultiplier tube (Hamamatsu) coupled to a 525/70 nm bandpass emission filter.

Imaging was performed in awake, embedded larvae expressing pan-neuronal nuclear GCaMP6s under the HuC/elavl3 promoter ^52^. Time-series data were acquired across nine planes spaced 25 µm apart using a piezoelectric objective scanner (PD72Z4CAA, Physik Instrumente), covering a total depth of 200 µm. The field of view (FOV) was sampled at 1024 ×512 pixels, images were acquired at 3 Hz for 100 seconds per trial. Each larva underwent two successive imaging sessions, one covering an anterior and one a posterior field of view, with the order of FOVs counterbalanced across fish. For each FOV, each of the 15 stimulus-concentration conditions was repeated three times (see above).

### ROI segmentation

Regions of interest were segmented from functional imaging data using Suite2p ^93^. Suite2p first performs rigid and non-rigid motion correction to compensate for sample drift and deformation during imaging. Frames were aligned to a reference image by maximizing cross-correlation. After motion correction, data were denoised and temporally filtered, and candidate regions of interest (ROIs) were detected based on local fluorescence correlations. Suite2p uses a combination of spatial clustering and neuropil correction to extract fluorescence traces from candidate neuronal regions. Following automated segmentation, ROIs were manually curated by visual inspection of both their spatial footprints and temporal activity traces to exclude artefacts or non-neuronal structures.

### Fluorescence preprocessing and neuronal filtering

For each neuron, fluorescence traces were extracted from segmented ROIs converted to ΔF/F_0_ using:

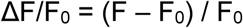

where F is the raw fluorescence signal and F_0_ is the mean fluorescence during the pre-stimulus baseline window. The baseline window corresponded to frames 80 - 150, approximately 23 s immediately preceding stimulus onset.

To standardize responses across neurons, traces were z-scored relative to the baseline window:

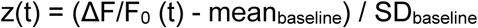

where mean_baseline_ and SD_baseline_ are the mean and standard deviation of the prestimulus baseline window. Neurons were filtered to exclude traces with insufficient signal quality before downstream analysis.

### Anatomical registration

Individual experiments were registered to a reference atlas coordinate system using ANTsPy, the Python interface for ANTs ^94^, together with custom-written code ^95^. At the end of each experiment, a high-resolution anatomical stack was acquired at 1 µm axial resolution. A manual affine transformation was first performed to align each anatomical stack with the Tg(*elavl3:Hsa.H2B-GCaMP6s*) reference stack from the mapzebrain atlas ^53^. This was done in Napari by visually matching corresponding anatomical landmarks ^96^. The resulting transformation was then used as an initialization for automated registration. Registered neuronal coordinates were assigned to anatomical regions using mapzebrain brain-area masks. The following abbreviations are used: aTriMN - anterior trigeminal motor nucleus; Cb - cerebellum; dHb - dorsal habenula; dTh - dorsal thalamus; EmT - eminentia thalami; fMN - facial motor nucleus; HC - caudal hypothalamus; HI - intermediate hypothalamus; HR - rostral hypothalamus; ifDMO - inferior dorsal medulla oblongata; ifVMO - inferior ventral medulla oblongata; IO - inferior olive; IPN - interpeduncular nucleus; iRaphe - inferior raphe; itDMO - intermediate dorsal medulla oblongata; itVMO - intermediate ventral medulla oblongata; MLR - mesencephalic locomotor region; NI - nucleus isthmi; nMLF - nucleus of the medial longitudinal fasciculus; OB - olfactory bulb; OE - olfactory epithelium; Pal - pallium; POA - preoptic area; PT - posterior tuberculum; PTec - pretectum; PTh - prethalamus; pTriMN - posterior trigeminal motor nucleus; sDMO - superior dorsal medulla oblongata; SGN - secondary gustatory nucleus; sRaphe - superior raphe; SubP - subpallium; svMO - superior ventral medulla oblongata; Tect - optic tectum; Teg - tegmentum; Tori - tori; vHb - ventral habenula; vMN - vagus motor nucleus; VSL - vagal sensory lobe

### Functional imaging analysis Averaged time-series visualization

To visualize average stimulus-evoked dynamics (**Fig. 3, Fig. S4**), neuronal response traces were averaged across trials, neurons, and fish separately for each stimulus. For each stimulus, fluorescence time series from all trials were aligned to stimulus onset. The mean and standard error of the mean were computed at each time point. These traces represented population-level stimulus responses and incorporated both inter-neuronal and inter-individual variability.

### Lifetime sparseness analysis

Lifetime sparseness was calculated to quantify how broadly or selectively (“sharply tuned”) individual neurons responded across stimuli. For each neuron, a response vector was computed from the mean responses to the seven chemosensory stimuli at their highest concentrations. Lifetime sparseness (S) was calculated as:

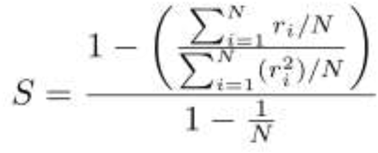

where (r_i_) is the mean response to stimulus *i*, and *N* is the number of stimuli. Values close to 0 indicate broad responses across stimuli, whereas values close to 1 indicate selective responses to a small number of stimuli.

### Response latency analysis

Response latency was estimated from regional mean fluorescence traces rather than from individual neurons. For each fish, stimulus condition, and anatomical region, fluorescence traces were averaged across all neurons assigned to that region. The resulting area-averaged trace was smoothed with a Gaussian filter (σ = 3 frames), and its first derivative was calculated. Latency was defined as the first frame after effective stimulus arrival at the larva, estimated to be at 53 s from trial start based on fluorescein calibration experiments, at which the derivative exceeded 0.01. Latencies were computed independently for each fish and region. Because this measure depends on the rise of the region-averaged fluorescence trace, latency estimates can be influenced by signal amplitude, response synchrony across neurons, and SNR within a region. Thus, differences between regions (e.g. between OE and OB) should be interpreted as latency differences in the averaged regional Ca^2+^ signal, rather than direct measurements of synaptic or sensory-transduction timing.

### Clustering analysis

For concentration-response clustering, analyses were restricted to the four stimuli presented at three concentrations (Fex, Pro, Cad, and Qun). For each stimulus separately, neurons significantly responsive to the highest concentration were selected, and a concentration-response vector was computed from the mean response amplitude at the low, intermediate, and high concentration.

Response vectors were MinMax-normalized to the range 0-1 and clustered using k-means clustering implemented in scikit-learn ^97^. The number of clusters was guided by silhouette analysis. Descriptive cluster labels were then assigned by inspecting the mean concentration-response profile of each cluster and matching recurring profiles across stimuli, yielding four response classes: ‘shallow-ramping’, ‘steep-ramping’, ‘high-threshold’, and ‘low-threshold’ neurons.

To test whether VENs within each valence class formed stimulus-specific subgroups, +VENs and −VENs were then clustered separately using k-means with k values from 2 to 5. Clustering was based on each neuron’s mean response amplitudes to the individual stimuli. Silhouette scores were low for both +VENs (max. (k=2): 0.33; min. (k=5): 0.23) and −VENs (max. (k=3): 0.26; min. (k=5): 0.17) indicating weak within-valence separability. For visualization, a common k value of 4 was chosen (+VENs, silhouette (k=4): 0.32; −VENs, silhouette (k=4): 0.22) was chosen as an intermediate, common cluster number. All clusters showed stronger responses to stimuli of their own valence than to stimuli of the opposite valence, and no clear anatomical segregation of clusters was observed.

### Regressors and correlation analyses

Neuronal responses to stimuli and behavior were analyzed using binary regressors encoding event timing. Although stimulus channels were switched for 10 s, dye measurements indicated slower wash-in and washout at the larva, with stimulus levels persisting for ≥ 20s (**Fig. S2D**). Stimulus-related regressors therefore used a 25 s window aligned to the estimated stimulus arrival, reflecting effective stimulus exposure rather than pump-command duration. Regressors were convolved with a double-exponential response kernel to approximate GCaMP6s fluorescence dynamics. :

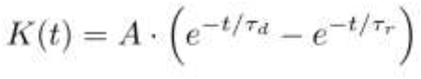

where *A* is an amplitude factor, (τ_r_) is the rise time constant (1.44 s), and (τ_d_) is the decay time constant (5 s), estimated from the indicator kinetics ^70^. Pearson’s correlation coefficient was then calculated between each neuron’s activity trace and the convolved regressor to quantify their association.

### Brain map visualization

Brain maps were generated to display the fraction of each neuronal response type within each brain region. Brain-area masks were obtained from the mapzebrain atlas ^53^ and applied to registered neuronal coordinates. To identify regions enriched for different neuron classes (e.g. stimulus-responsive, behaviorally correlated etc.), neurons were first assigned to anatomical regions using mapzebrain masks. For each region, we counted the number of neurons fitting the respective criterion and compared this count to the number expected based on the fish-specific fraction of stimulus-responsive neurons across all recorded neurons using a one-sided binomial test. P-values were combined across animals using Fisher’s method. Regions with significantly higher fractions than expected were highlighted in the brain maps (p < 0.05).

### Identification of stimulus-responding neurons

To identify neurons that responded consistently to stimuli, regression was first performed using the three repeated trials of each stimulus at each concentration. This analysis identified neurons with reliable responses across repeated presentations of the same stimulus. For the brain maps shown in **Fig. 3**, we used a threshold of *r* > 0.3 to any highest-concentration stimulus regressor. Results of other thresholding are shown in **Fig. S3**.

### Identification of stimulus-specific neurons

Stimulus-specific regressors were constructed to identify neurons responding selectively to individual stimuli. This analysis was restricted to the highest-concentration trials, yielding 21 trials in total: three presentations for each of the seven stimuli. Each regressor contained non-zero values only during presentations of the stimulus of interest, allowing neurons preferentially responding to a single stimulus to be distinguished from neurons with broader multi-stimulus tuning.

### Identification of valence-selective neurons

Valence regressors were constructed to identify neurons whose activity was aligned with stimulus groups classified behaviorally as appetitive or aversive (**Figs. 1,2**). HCl was excluded to maintain balanced stimulus categories, resulting in valence groups that each contained two stimuli typically considered to act primarily as odorants (Ade, Cad; Fex, Kin) and one typically considered to act primarily as tastant (Qun; Pro).

Separate regressors encoded appetitive and aversive stimulus presentations, and neurons significantly correlated with these regressors were classified as positive or negative valence-encoding neurons (+VENs or −VENs), respectively. VENs were therefore defined operationally by shared responses across multiple stimuli associated with the same behavioral output, rather than by selectivity for individual chemical identities. However, this definition does not by itself fully distinguish valence-related activity from other co-aligned features, including shared motor-state changes

### Behavior-related neuronal activity

Behavior-related neuronal activity was examined by regressing neuronal fluorescence traces against behavioral traces derived from angular tail movements.This analysis identified neurons whose activity correlated with motor output rather than stimulus presentation (behavior-correlated neurons: “BCNs”). Because these regressors were constructed from behavioral traces across the entire trial, BCNs were interpreted as neurons with activity correlated with movement-related variables, rather than as neurons specifically or exclusively driven by chemosensory-evoked motor output. To capture activity associated with left- or right-biased movements, lateralized behavioral regressors were constructed. Turning bouts, defined as swim events with measurable lateralized tail angles (see above), were detected and binarized into a trace indicating whether the fish was turning (value of 1) or not (value of 0). This binary turning trace was then convolved with the GCaMP kernel to generate a regressor reflecting the expected neural dynamics associated with turning behavior, regardless of direction.

These regressors were used to identify neurons whose activity was correlated with lateralized swim events and turning-related behavior (turn-BCNs). Similarly, a straight bout regressor was constructed following the same procedure to identify neurons whose activity was correlated with straight swim events (straight-BCNs). In addition to the turning and straight bout regressors, a tail vigor regressor was calculated from tail-angle variability (see above) and convolved with the same GCaMP kernel used for stimulus regressors to identify neurons whose activity was correlated with general behavioral vigor.

### Statistical significance of regression analyses

Statistical significance for stimulus-specific, valence, and behavior regressors (see above) was assessed using permutation testing. To generate null distributions, trial assignments were randomly shuffled 1,000 times, disrupting the temporal relationship between neuronal activity and stimulus or behavior. After each shuffle, Pearson’s correlation was recalculated between the neuronal activity trace and the permuted regressor. The p-value was calculated as the fraction of shuffled correlations exceeding the observed correlation. False discovery rate (FDR) correction (Benjamini–Hochberg procedure) was applied to all neurons within each fish, controlling the expected proportion of false positives at < 0.05 (see below). Neurons with FDR-corrected p_FDR_ < 0.05 were classified as significantly correlated.

### Population response vectors

Population response vectors were used to quantify stimulus-evoked activity within defined temporal response windows. Unless stated otherwise, responses were quantified in a 20 s window starting at stimulus onset, providing a comprehensive measure of stimulus-evoked activity that captured both early response onset and later response phases. For each fish, a grand average response was computed by averaging neuronal activity across all recorded neurons during stimulus presentation. The peak of this response was identified, and the time at which activity reached 50% of the peak was used to dynamically align response quantification across experiments. This approach allowed response windows to reflect the stimulus-evoked dynamics of each experiment rather than relying exclusively on fixed time intervals.

### Population decoding of stimulus identity

Population decoding was used to assess how well neuronal population activity distinguished between stimuli. Template matching was used because it is robust to small sample sizes and does not require strong assumptions about data distributions. For each fish, neurons were grouped by anatomical brain area and population response vectors were constructed (as described above). Only brain areas containing at least five neurons were included in the analysis. Classification analyses were performed using trials corresponding to the highest available concentration for each stimulus that was presented in a concentration series. For stimulus identity classification, seven stimuli served as labels. Within each brain area and fish, trials were split into training and test sets using stratified k-fold cross-validation. Three-fold cross-validation was used for stimulus identity decoding. Training data were used to compute mean population vectors, or templates, for each class (stimulus). Test trials were classified by comparing their population vectors with the templates using cosine distance. Each test trial was assigned to the class with the smallest cosine distance, and classification accuracy was averaged across folds to obtain the mean score for each brain area and fish.

### Permutation testing for decoding analyses

To determine whether classification accuracy exceeded chance, trial labels were randomly shuffled 1,000 times, and the full classification procedure was repeated for each shuffled dataset. The p-value was calculated as the fraction of permutations with classification accuracy exceeding the observed value. P-values across fish were combined for each brain area using Fisher’s method, followed by FDR correction for multiple comparisons. Brain areas with FDR-corrected p < 0.05 were considered to encode the decoded variable at the population level. For each brain area, mean and standard deviation of classification accuracy across fish were reported. Confusion matrices (for stimulus identity, averaged across fish) and accuracy distributions were visualized to illustrate decoding performance. In addition, classification accuracies were projected onto a brain map, analogous to the correlation analysis, to visualize the spatial distribution of decoding performance across anatomical regions. Areas with significant classification (FDR-corrected p < 0.05) were highlighted on these maps to emphasize regions that showed encoding of stimulus identity.

### Spatial distribution analysis

Kernel density estimation (KDE) was used to visualize and quantify the spatial distributions of positive and negative VENs along the medial-lateral, anterior-posterior, and dorsal-ventral axes. KDEs were estimated separately for each valence group using Scott’s rule for bandwidth selection (a standard method that scales bandwidth with sample size), ensuring consistent smoothing across comparisons. The evaluation range was restricted to the observed neuronal coordinates within each brain region.

The degree of overlap between positive and negative VEN distributions was quantified from the KDEs using the overlap integral. Spatial separation between groups was further assessed using linear discriminant analysis. For each region, VEN coordinates were used to train an LDA classifier to discriminate positive from negative VENs. For each brain area, neuronal x, y, and z coordinates and valence labels were used as input to LDA. This yielded a one-dimensional projection of each neuron and a separation vector, whose angle relative to the anatomical axes was used to assess alignment with anatomical directions. Separation strength was quantified using the Fisher ratio, 5-fold cross-validated classification accuracy, and area under the ROC curve. For visualization, neurons were plotted in 3D together with the LDA vector and its orthogonal separating plane, and one-dimensional LDA projections were displayed using kernel density estimation.

### Behavior modulation analysis Classification of trial behavior strength

Trials were categorized according to stimulus-evoked changes in swimming activity. The number of swim bouts was counted in two temporal windows relative to stimulus onset: a 20 s pre-stimulus window and a 20 s stimulus window. For each trial, the ratio of post-stimulus to pre-stimulus bout counts was computed. Trials were classified into behavioral categories using a 10% change threshold. Trials with increased bout counts were classified as movement-increased trials, trials with decreased bout counts as movement-reduced trials, and trials with changes below threshold as unchanged. The threshold of 10% change was chosen to achieve a comparable number of trials in each category. Analyses included 90 POS trials and 120 NEG trials across the population (3 or 4 stimuli × 10 fish × 3 trials).

### Behavioral modulation index of neural responses

A behavioral modulation index was calculated to quantify how strongly neuronal responses were modulated by behavioral state (**Fig. 7**). Trials were first classified into movement-increased and movement-decreased classes (see above). For each class, fluorescence traces were averaged within each brain area, pooling across all neurons. Only areas containing at least ten neurons were included to ensure robustness. Response amplitudes were then calculated for each behavioral condition, and modulation was quantified from the difference between strong- and weak-behavior trial responses:

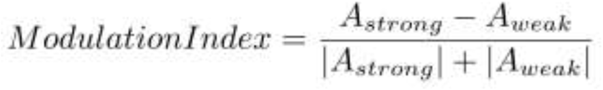

where A_strong_ and A_weak_ are the response amplitudes during movement-increased and movement-decreased trials, respectively. This index ranges from –1 to 1, with positive values indicating stronger responses during movement-increased trials, and negative values indicating stronger responses during movement-decreased trials.

### Pairwise neuronal correlation analyses

Activity co-fluctuations between neuronal populations, sometimes referred to as functional “coupling”, were quantified by calculating pairwise correlations during baseline and stimulus periods. These are referred to here as spontaneous and noise correlations, respectively. Correlation analyses were performed within individual fish, and resulting metrics were subsequently averaged across fish. Brain area pairs were included in the analysis only if the total number of pairwise correlations pooled across fish exceeded 100. Neurons were assigned to functional groups based on prior classification analyses, specifically into valence-encoding neurons (VENs) and behaviorally correlated neurons (BCNs). Correlations were computed within a group (e.g., among +VENs) as well as between groups (e.g., between +VENs in area A and BCNs in area B). For each pair of brain areas, all pairwise Pearson correlation coefficients were calculated for the specified groups, and their distributions were recorded for each fish.

### Spontaneous and noise correlation analysis

Spontaneous correlations were used to quantify correlated activity during baseline periods. For each neuron, activity during a 23 s pre-stimulus baseline period was extracted for each trial (total of 45 trials) and concatenated to generate baseline activity vectors. Pearson correlation coefficients were then computed between baseline activity vectors for neuron pairs within and between brain areas, separately for each fish.

Noise correlations were used to quantify trial-to-trial co-variability after removing stimulus-driven responses. For each neuron, response amplitude was defined as the mean z-scored activity within the stimulus response window, yielding one response value per trial. To isolate residual activity, the mean response across trials of the same stimulus was subtracted from each trial-wise response amplitude. Pearson correlation coefficients were then computed between residual response vectors for neuron pairs.

For both spontaneous and noise correlations, correlation distributions for each area pair were summarized using the Fisher z-transformed mean correlation. Correlation statistics were first computed separately for each fish and then combined across animals.

Statistical significance was assessed using null sampling distributions. For each area pair, null distributions were generated by repeatedly computing correlations from randomly selected neurons drawn from the same brain areas and matched in number to the neuron populations of interest. This procedure was repeated 1,000 times. Empirical p-values were calculated by comparing observed correlation statistics to the corresponding null distributions. P-values were combined across animals using Fisher’s method and corrected for multiple comparisons across area pairs using false discovery rate correction.

### Software and statistical analysis

All primary data analysis was performed in Python 3.9 using scientific computing libraries including NumPy, SciPy, pandas, and scikit-learn. ROI segmentation and motion correction of calcium imaging data were performed using Suite2p. Image registration and brain-area mapping were performed using ANTsPy and Napari, with alignment to the mapzebrain atlas ^53^.

Statistical analyses used nonparametric tests where appropriate. Wilcoxon-Mann-Whitney U tests were used for unpaired comparisons, and Wilcoxon signed-rank tests were used for paired comparisons. A Dunn-Hollander-Wolfe test was used for nonparametric multiple comparisons with one control group (**Fig. 1E**), where data was not paired (reported p values are adjusted for multiple comparisons). Linear mixed-effects models were implemented using the statsmodels package in Python and were used to analyze stimulus-evoked changes in bout metrics while accounting for repeated measurements across animals and imaging sessions. Multiple testing correction was performed using the Benjamini-Hochberg false discovery rate (FDR) procedure (q = 0.05).

Figures were initially generated using Matplotlib and Seaborn, with three-dimensional visualizations implemented using Matplotlib’s mplot3d and custom code for rendering vectors and planes, and finally assembled in Illustrator (Adobe). Computationally intensive analyses, including permutation testing and pairwise correlation analyses, were parallelized where possible. All analyses were implemented in custom Python scripts, version-controlled with Git, and random processes were seeded to ensure reproducibility.

## Detailed statistics

**Figure 1E:** change in stimulus zone occupancy vs. control experiments differed for Ade (*p* = 0.00080), Cad (*p* = 0.00024), HCl (*p* = 8.0 × 10⁻^7^), Qun (*p* = 3.2 × 10⁻^12^), Fex (p = 0.00016), Kin (p = 1.8 × 10⁻^5^), and Pro (*p* = 0.015); Dunn-Hollander-Wolfe test vs. control.

Figure 1F: Stimulus-zone occupancy at the individual-experiment level differed from control experiments for NEG (*p* = 3.1 × 10⁻^6^) and POS (*p* = 0.00025); Wilcoxon-Mann-Whitney test.

Figure 1H: Change in heading angle differed between NEG and POS (*p* = 0.00062); Wilcoxon-Mann-Whitney test.

Figure 1I: Swim speed within the stimulus zone compared with the corresponding no-stimulus zone differed in NEG (*p* = 5.8 × 10⁻^6^) and POS (*p* = 0.0025); Wilcoxon signed-rank test.

Figure 1J: Experiment-wise swim-speed change relative to the no-stimulus differed between NEG and POS stimuli (*p* = 7.9 × 10⁻^8^); Wilcoxon-Mann-Whitney test.

Figure 1K: Swim speed within the two zones did not differ in Ctrl (*p* = 0.81); Wilcoxon signed-rank test.

Figure 2E: Posture angle differed between Qun 25 µM and Qun 2.5 mM (*p* = 2.83 × 10⁻²⁸) and between Fex 1:1000 and Fex 1:10 (*p* = 1.81 × 10⁻¹⁰); linear mixed-effects model (LMM).

Figure 2F: Posture angle differed between pre- and post-stimulus periods for Ade (*p* = 7.81 × 10⁻¹⁴), Cad (*p* = 4.76 × 10⁻¹⁷), HCl (*p* = 4.15 × 10⁻¹⁰), Qun (*p* = 2.92 × 10⁻³⁶), Fex (*p* = 3.23 × 10⁻²⁴), Kin (*p* = 2.38 × 10⁻²⁷), and Pro (*p* = 3.79 × 10⁻¹⁷); LMM.

Figure 2G: Bout count differed from baseline in the 6–9 s (*p* = 0.0195), 9–12 s (*p* = 0.0195), 12–15 s (*p* = 0.0273), 15–18 s (*p* = 0.0059), 18–21 s (*p* = 0.002), 21–24 s (*p* = 0.002), 24–27 s (*p* = 0.0273), and 27–30 s (*p* = 0.0137) time bins for negative stimuli, and in the 3–6 s (*p* = 0.024), 15–18 s (*p* = 0.037) time bins for positive stimuli; Wilcoxon signed-rank test.

Figure 2I: Turning bout frequency differed between pre- and post-stimulus periods for negative stimuli (*p* = 0.0273) but not positive stimuli (*p* = 0.1387); Wilcoxon signed-rank test.

Figure 2J: Turning bout frequency differed between pre- and post-stimulus periods for Ade (*p* = 0.0103), Cad (*p* = 0.0339), HCl (*p* =0.0344), and Qun (*p* = 0.0374), but not for Fex (*p* = 0.2989), Kin (*p* = 0.4183), or Pro (*p* = 0.1584); LMM.

Figure 3F: Response latency differed between forebrain and hindbrain neurons (*p* = 0.0137); Wilcoxon signed-rank test.

Figure 3M: Brain areas enriched with stimulus-responsive neurons included OB (*p* < 1 × 10⁻²⁵⁵), OE (*p* < 1 × 10⁻²⁵⁵), Pal (*p* < 1 × 10⁻²⁵⁵), SubP (*p* < 1 × 10⁻²⁵⁵), ifDMO (*p* = 7.30 × 10⁻¹⁶⁰), sDMO (*p* = 3.06 × 10⁻¹⁸¹), POA (*p* = 5.08 × 10⁻¹⁹⁵), EmT (*p* = 3.59 × 10⁻¹⁰²), itVMO (*p* = 2.96 × 10⁻⁸⁶), NI (*p* = 6.08 × 10⁻⁷⁶), fMN (*p* = 2.69 × 10⁻⁴²), itDMO (*p* = 1.46 × 10⁻⁴⁵), PTh (*p* = 3.64 × 10⁻²²), vHb (*p* = 1.61 × 10⁻³²), sVMO (*p* = 6.22 × 10⁻²⁹), vMN (*p* = 7.51 × 10⁻¹⁵), IO (*p* = 2.17 × 10⁻⁸), LC (*p* = 7.33 × 10⁻⁸), aTriMN (*p* = 4.02 × 10⁻⁶), ifVMO (*p* = 1.31 × 10⁻⁵), IPN (*p* = 6.15 × 10⁻⁵), iRaphe (*p* = 0.037), VSL (*p* = 0.014), and HI (*p* = 0.012); one-sided binomial tests with p-values combined across fish using Fisher’s method.

Figure 4A: Stimulus classification accuracy was significantly above chance in dHb (*p* < 1 × 10⁻²⁵⁵), IPN (*p* < 1 × 10⁻²⁵⁵), sRaphe (*p* < 1 × 10⁻²⁵⁵), LC (*p* < 1 × 10⁻²⁵⁵), Cb (*p* < 1 × 10⁻²⁵⁵), HR (*p* < 1 × 10⁻²⁵⁵), HI (*p* < 1 × 10⁻²⁵⁵), PT (*p* < 1 × 10⁻²⁵⁵), POA (*p* < 1 × 10⁻²⁵⁵), PTh (*p* < 1 × 10⁻²⁵⁵), SubP (*p* < 1 × 10⁻²⁵⁵), OB (*p* < 1 × 10⁻²⁵⁵), OE (*p* < 1 × 10⁻²⁵⁵), EmT (*p* < 1 × 10⁻²⁵⁵), Pal (*p* < 1 × 10⁻²⁵⁵), vHb (*p* < 1 × 10⁻²⁵⁵), SGN (*p* < 1 × 10⁻²⁵⁵), sDMO (*p* < 1 × 10⁻²⁵⁵), PTec (*p* < 1 × 10⁻²⁵⁵), dTh (*p* < 1 × 10⁻²⁵⁵), NI (*p* = 1.6 × 10⁻⁵), sVMO (*p* = 4.8 × 10⁻⁵), Tori (*p* = 6.3 × 10⁻⁵), iDMO (*p* = 4.3 × 10⁻⁴), aTriMN (*p* = 4.3 × 10⁻⁴), Teg (*p* = 4.3 × 10⁻⁴), iVMO (*p* = 1.9 × 10⁻³), iDMO (*p* = 6.6 × 10⁻³), IO (*p* = 6.6 × 10⁻³), HC (*p* = 8.8 × 10⁻³), and iRaphe (*p* = 0.044); permutation testing with p-values combined across fish using Fisher’s method.

Figure 4C: Decoding accuracy differed between OB and OE (*p* = 0.0039), SubP (*p* = 0.0039), POA (*p* = 0.0020), EmT (*p* = 0.0049), dHb (*p* = 0.0010), vHb (*p* = 0.0010), PTh (*p* = 0.0020), HR (*p* = 0.0010), and HI (*p* = 0.0010), but not between OB and Pal (*p* = 0.1563); Wilcoxon signed-rank test.

Figure 5F: Areas enriched in valence-encoding neurons included SubP (*p* = 4.94 × 10⁻⁷¹), Pal (*p* = 1.07 × 10⁻⁴⁴), POA (*p* = 1.73 × 10⁻²⁶), OB (*p* = 1.39 × 10⁻¹⁶), OE (*p* = 1.46 × 10⁻⁸), PT (*p* = 1.12 × 10⁻⁴), vHb (*p* = 5.25 × 10⁻⁵), ifDMO (*p* = 4.11 × 10⁻⁵), HI (*p* = 3.87 × 10⁻³), and sDMO (*p* = 0.035); one-sided binomial tests with p-values combined across fish using Fisher’s method.

Figure 5G: Spatial separation between positive and negative valence-encoding neurons was observed in OB (AUC = 0.70 ± 0.06, *p* = 9.99 × 10⁻⁴), SubP (AUC = 0.72 ± 0.02, *p* = 9.99 × 10⁻⁴), POA (AUC = 0.66 ± 0.15, *p* = 0.013), PT (AUC = 0.82 ± 0.10, *p* = 9.99 × 10⁻⁴), and HI (AUC = 0.72 ± 0.14, *p* = 0.023), but not in Pal (AUC = 0.57 ± 0.06, *p* = 0.056); LDA classification with label-shuffled permutation testing (1,000 permutations).

Figure 5I: Mean noise correlations were greater than expected by chance for −VEN pairs (*p* < 0.001) and +VEN pairs (*p* < 0.001), and lower than expected by chance for −VEN/+VEN pairs (*p* < 0.001); comparison against null sampling distributions generated from randomly sampled neurons (1,000 permutations); Wilcoxon–Mann–Whitney test.

Figure 5J: Significant positive noise correlations (FDR-corrected *p* < 0.05) were observed between multiple area pairs within the −VEN and +VEN populations, whereas significant negative noise correlations (FDR-corrected *p* < 0.05) were observed between multiple area pairs of the −VEN and +VEN populations; significance was determined by comparison against null sampling distributions generated from randomly sampled neurons (1,000 permutations).

Figure 5L: Mean spontaneous correlations were greater than expected by chance for −VEN pairs (*p* < 0.001) and +VEN pairs (*p* < 0.001), and lower than expected by chance for −VEN/+VEN pairs (*p* < 0.001); comparison against null sampling distributions generated from randomly sampled neurons (1,000 permutations); Wilcoxon–Mann–Whitney test.

Figure 5M: Significant positive spontaneous correlations (FDR-corrected *p* < 0.05) were observed between multiple area pairs within the −VEN and +VEN populations, whereas significant negative spontaneous correlations (FDR-corrected *p* < 0.05) were observed between multiple area pairs of the −VEN and +VEN populations; significance was determined by comparison against null sampling distributions generated from randomly sampled neurons (1,000 permutations).

**Figure 6C:** Areas enriched in turning behaviour-correlated neurons included Teg (*p* = 1.51 × 10⁻¹⁷⁶), itVMO (*p* = 1.07 × 10⁻¹⁷⁴), nMLF (*p* = 2.82 × 10⁻¹²²), ifDMO (*p* < 1 × 10⁻²⁵⁵), fMN (*p* = 9.40 × 10⁻¹⁰¹), NI (*p* = 8.89 × 10⁻¹⁰¹), PT (*p* = 1.81 × 10⁻⁸⁵), itDMO (p = 5.83 × 10⁻⁵⁸), sVMO (*p* = 9.76 × 10⁻²⁹), vMN (*p* = 1.39 × 10⁻¹⁹), POA (*p* = 3.75 × 10⁻¹³), ifVMO (*p* = 5.90 × 10⁻¹²), sDMO (p = 1.27 × 10⁻¹⁰), LC (*p* = 1.09 × 10⁻¹⁰), dTh (*p* = 1.62 × 10⁻⁹), iRaphe (*p* = 6.67 × 10⁻³), VSL (*p* = 6.00 × 10⁻⁶), and PTec (*p* = 5.00 × 10⁻⁵); one-sided binomial tests performed per fish and combined across animals using Fisher’s method.

Figure 7D: Mean response amplitudes of −VENs differed between weak and strong behavioural trials for positive stimuli (*p* = 0.034) and negative stimuli (*p* = 0.0022); Wilcoxon–Mann–Whitney Test.

Figure 7E: Mean response amplitudes of +VENs did not differ between weak and strong behavioural trials for positive stimuli (*p* = 0.685) or negative stimuli (*p* = 0.462); Wilcoxon–Mann–Whitney Test.

Figure 7H: Correlations with the turning regressor differed between −VENs and +VENs (*p* = 3.25 × 10⁻¹⁷⁸); Wilcoxon–Mann–Whitney test.

Figure 7I: Correlations between Turn-BCNs and the negative versus positive valence regressors differed significantly (*p* < 1 × 10⁻²⁵⁵); Wilcoxon signed-rank test.

Figure 7M: Mean stimulus-evoked noise correlations were greater than expected by chance for −VEN/BCN pairs (*p* < 0.001) and +VEN/BCN pairs (*p* < 0.001), comparison against matched-number randomly sampled neurons; Wilcoxon–Mann–Whitney test.

Figure 7N: Significant positive stimulus-evoked noise correlations (FDR-corrected *p* < 0.05) were observed between multiple area pairs of −VENs and Turn-BCNs, whereas significant negative stimulus-evoked noise correlations (FDR-corrected *p* < 0.05) were observed between multiple area pairs of +VENs and Turn-BCNs; significance was determined by comparison against null sampling distributions generated from randomly sampled neurons (1,000 permutations).

Figure 7O: Mean spontaneous correlations were greater than expected by chance for −VEN/BCN pairs (*p* < 0.001) and lower than expected by chance for +VEN/BCN pairs (*p* < 0.001), comparison against matched-number randomly sampled neurons; Wilcoxon–Mann–Whitney test.

Figure 7P: Significant positive spontaneous correlations (FDR-corrected *p* < 0.05) were observed between multiple area pairs of −VENs and Turn-BCNs, whereas significant negative spontaneous correlations (FDR-corrected *p* < 0.05) were observed between multiple area pairs of +VENs and Turn-BCNs; significance was determined by comparison against null sampling distributions generated from randomly sampled neurons (1,000 permutations).

Figure 7Q: Stimulus-evoked noise correlation distributions differed between sparse/BCN pairs and matched-number randomly sampled neurons (*p* < 0.001); Wilcoxon–Mann–Whitney test.

Figure 7R: Significant stimulus-evoked noise correlations (FDR-corrected *p* < 0.05) were observed between multiple area pairs of sparse neurons and Turn-BCNs; significance was determined by comparison against null sampling distributions generated from randomly sampled neurons (1,000 permutations).

Figure 7S: Spontaneous correlation distributions for sparse/BCN pairs differed from those of matched-number randomly sampled neurons (*p* < 0.001); Wilcoxon–Mann–Whitney test.

Figure 7T: Significant spontaneous correlations (FDR-corrected *p* < 0.05) were observed between multiple area pairs of sparse neurons and Turn-BCNs; significance was determined by comparison against null sampling distributions generated from randomly sampled neurons (1,000 permutations).

**Supplementary Figure S1F**: Change in heading angle differed between NEG and POS for individual larvae (*p* = 8.6 × 10⁻¹¹); Wilcoxon-Mann-Whitney test.

**Supplementary Figure S1G**: Swim speed within the stimulus zone compared with the corresponding no-stimulus zone differed in both NEG (*p* = 4.0 × 10⁻¹⁵) and POS (*p* = 2.1 × 10⁻¹²) for individual larvae; Wilcoxon signed-rank test.

**Supplementary Figure S1H**: Larva-wise swim-speed change relative to the no-stimulus differed between NEG and POS stimuli(*p* = 7.9 × 10⁻^8^); Wilcoxon-Mann-Whitney test.

**Supplementary Figure S1I**: Swim speed within the two zones did not differ in Ctrl (*p* = 0.93); Wilcoxon signed-rank test.

**Supplementary Figure S2A:** Posture angle differed between Cad 25 μM and Cad 2.5 mM (*p* = 1.20 × 10⁻⁷); linear mixed-effects model (LMM).

**Supplementary Figure S2B:** Posture angle differed between Pro 25 μM and Pro 2.5 mM (*p* = 1.86 × 10⁻¹⁰); LMM.

**Supplementary Figure S2C:** For Qun (2.5 mM), posture angle differed between pre- and stimulus periods for left-directed stimulus delivery (45°; *p* = 1.69 × 10⁻⁷) and right-directed stimulus delivery (−45°; *p* = 8.71 × 10⁻⁵), but not for center-directed stimulus delivery (0°; *p* = 0.102). For Pro (2.5 mM), posture angle differed between pre- and stimulus periods for left-directed stimulus delivery (45°; *p* = 0.0459) and right-directed stimulus delivery (−45°; *p* = 0.0400), but not for center-directed stimulus delivery (0°; *p* = 0.217); LMM.

**Supplementary Figure S2F:** Turning bout frequency differed from baseline in the 3–6 s (*p* = 0.0488), 15–18 s (*p* = 0.0273) and 27–30 s (*p* = 0.0488) time bins for positive stimuli. For negative stimuli, turning bout frequency differed from baseline in the 6–9 s (*p* = 0.0195), 9–12 s (*p* = 0.0273), 12–15 s (*p* = 0.0273), 15–18 s (*p* = 0.0195), 18–21 s (*p* = 0.0195), and 21–24 s (*p* = 0.0137) time bins. Straight bout frequency did not differ from baseline for positive stimuli. For negative stimuli, straight bout frequency differed from baseline in the 15–18 s (*p* = 0.0137), 18–21 s (*p* = 0.0273), and 21–24 s (*p* = 0.0371) time bins. Straight bout frequency did not differ between pre- and post-stimulus periods for negative stimuli (*p* = 0.922) or positive stimuli (*p* = 0.846); Wilcoxon signed-rank test.

**Supplementary Figure S2G:** Turning bout frequency did not differ between pre- and post-stimulus periods for Cad (250 μM; *p* =0.942), Qun (250 μM; *p* = 0.174), Fex (1:100; *p* = 0.636), Pro (250 μM; *p* = 0.420), Cad (25 μM; *p* = 0.847), Qun (25 μM; *p* = 0.253), Fex (1:1000; *p* = 0.462), or Pro (25 μM; *p* = 0.388); LMM.

**Supplementary Figure S3E:** For neurons responding to at least one stimulus with r ≥ 0.2, areas enriched in stimulus-responsive neurons included OE (*p* = 1.67 × 10⁻²⁶⁹), OB (*p* < 1 × 10⁻²⁵⁵), Pal (*p* < 1 × 10⁻²⁵⁵), sDMO (*p* = 5.90 × 10⁻²⁰⁹), POA (*p* = 1.58 × 10⁻¹⁴⁸), ifDMO (*p* = 4.65 × 10⁻¹⁴³), NI (*p* = 7.66 × 10⁻⁹⁵), itVMO (*p* = 7.51 × 10⁻⁸³), EmT (*p* = 7.74 × 10⁻⁷¹), itDMO (*p* = 7.01 × 10⁻⁶⁵), fMN (*p* = 2.45 × 10⁻⁴³), sVMO (*p* = 1.46 × 10⁻²⁹), vMN (*p* = 1.51 × 10⁻²⁴), vHb (*p* = 5.14 × 10⁻²⁰), PTh (*p* = 1.50 × 10⁻¹¹), IO (*p* = 1.38 × 10⁻¹¹), LC (*p* = 5.35 × 10⁻¹¹), aTriMN (*p* = 1.85 × 10⁻⁸), ifVMO (*p* = 2.11 × 10⁻⁵), VSL (*p* = 5.78 × 10⁻⁵), HI (*p* = 6.83 × 10⁻⁴), SGN (*p* = 5.09 × 10⁻³), and iRaphe (*p* = 0.0108). Using a stricter response threshold (r ≥ 0.5), enriched areas included OB (p < 1 × 10⁻²⁵⁵), Pal (*p* < 1 × 10⁻²⁵⁵), POA (*p* = 1.03 × 10⁻¹⁸⁵), EmT (*p* = 4.97 × 10⁻¹⁰⁶), OE (*p* = 5.44 × 10⁻⁶³), ifDMO (*p* = 1.55 × 10⁻⁴⁶), sDMO (*p* = 1.44 × 10⁻⁴⁴), PTh (*p* = 6.12 × 10⁻³⁶), itVMO (*p* = 1.47 × 10⁻²⁸), vHb (*p* = 1.30 × 10⁻²⁵), NI (*p* = 1.04 × 10⁻¹⁶), sVMO (*p* = 1.50 × 10⁻¹⁶), itDMO (*p* = 1.09 × 10⁻¹⁴), fMN (*p* = 4.94 × 10⁻⁸), LC (*p* = 2.69 × 10⁻³), IPN (*p* = 2.31 × 10⁻³), dHb (*p* = 8.74 × 10⁻⁴), and IO (*p* = 0.0461); one-sided binomial tests performed per fish and combined across animals using Fisher’s method.

**Supplementary Figure S3F**: Areas enriched in neurons responding to more than one stimulus (r ≥ 0.3) included POA (*p* = 1.85 × 10⁻²⁰⁸), ifDMO (*p* = 1.57 × 10⁻¹²²), sDMO (*p* = 1.88 × 10⁻¹¹¹), OE (*p* = 1.34 × 10⁻⁷⁴), itVMO (*p* = 1.12 × 10⁻⁷²), EmT (*p* = 2.83 × 10⁻⁷⁰), NI (*p* = 4.06 × 10⁻⁵⁵), OB (*p* < 1 × 10⁻²⁵⁵), Pal (*p* < 1 × 10⁻²⁵⁵), fMN (*p* = 2.37 × 10⁻³²), vHb (*p* = 3.96 × 10⁻²⁶), sVMO (*p* = 2.53 × 10⁻²⁵), itDMO (*p* = 1.50 × 10⁻¹⁶), PTh (*p* = 3.87 × 10⁻¹⁴), LC (*p* = 2.82 × 10⁻⁹), IPN (*p* = 2.49 × 10⁻⁶), ifVMO (*p* = 3.19 × 10⁻⁶), IO (*p* = 4.23 × 10⁻⁵), aTriMN (*p* = 3.91 × 10⁻³), vMN (*p* = 1.98 × 10⁻³), HI (*p* = 8.39 × 10⁻⁴), iRaphe (*p* = 0.0101), and dHb (*p* = 0.0294); one-sided binomial tests performed per fish and combined across animals using Fisher’s method.

**Supplementary Figure S4C**: Decoding accuracy was lower than in OB for all hindbrain areas shown (PTec, Tect, NI, Cb, SGN, sDMO, sRaphe, sVMO, aTriMN, itDMO, itVMO, iRaphe, fMN, ifDMO, ifVMO, vMN, and VSL; all *p* = 9.77 × 10⁻⁴); Wilcoxon signed-rank test, reflecting a consistent reduction in decoding performance across all animals.

**Supplementary Figure S4D:** Stimulus response similarity differed from OB for OE (*p* = 1.95 × 10⁻³), SubP (*p* = 5.86 × 10⁻³), Pal (*p* = 9.77 × 10⁻³), POA (*p* = 0.0195), HR (*p* = 0.0195), dHb (*p* = 1.95 × 10⁻³), and vHb (*p* = 0.0488); Wilcoxon signed-rank test.

**Supplementary Figure S4E:** Trial-to-trial reliability differed from OB for OE (*p* = 1.95 × 10⁻³), HI (*p* = 1.95 × 10⁻³), HR (*p* = 1.95 × 10⁻³), PT (*p* = 1.95 × 10⁻³), POA (*p* = 9.77 × 10⁻³), PTh (*p* = 9.77 × 10⁻³), dHb (*p* = 3.91 × 10⁻³), and vHb (*p* = 3.91 × 10⁻³), but did not differ from OB for EmT (*p* = 0.0840), Pal (*p* = 0.160), or SubP (*p* = 0.770); Wilcoxon signed-rank test.

**Supplementary Figure S4F:** Response latency was longer than in OB for OE (*p* = 1.95 × 10⁻³), PT (*p* = 1.95 × 10⁻³), HR (*p* = 1.95 × 10⁻³), PTh (*p* = 5.86 × 10⁻³), vHb (*p* = 7.81 × 10⁻³), and HI (*p* = 0.0156), but did not differ from OB for Pal (*p* = 0.688), SubP (*p* = 0.188), POA (*p* = 0.195), EmT (*p* = 0.625), or dHb (*p* = 0.0938); Wilcoxon signed-rank test.

**Supplementary Figure S4G:** Lifetime sparseness was greater than in OB for OE (*p* = 3.91 × 10⁻³), indicating more selective responses to individual stimuli. Lifetime sparseness was lower than in OB for HI (*p* = 1.95 × 10⁻³), HR (*p* = 1.95 × 10⁻³), POA (*p* = 1.95 × 10⁻³), PT (*p* = 1.95 × 10⁻³), SubP (*p* = 1.95 × 10⁻³), dHb (*p* = 1.95 × 10⁻³), vHb (*p* = 1.95 × 10⁻³), and Pal (*p* = 5.86 × 10⁻³), indicating broader responses across multiple stimuli. Lifetime sparseness did not differ from OB for EmT (*p* = 0.322) or PTh (*p* = 0.0645); Wilcoxon signed-rank test.

**Supplementary Figure** S**5D:** The spatial distributions of positive and negative VENs differed along the dorsal–ventral axis in OB (*p* = 1 × 10⁻²⁵⁵), SubP (*p* = 1 × 10⁻²⁵⁵), POA (*p* = 1 × 10⁻²⁵⁵), and PT (*p* = 1 × 10⁻²⁵⁵); along the lateral–medial axis in POA (*p* = 0.0148); and along the anterior–posterior axis in Pal (*p* = 0.0438), PT (*p* = 0.0007), and HI (*p* =0.0026). No significant differences were detected along the remaining axes shown; two-sample Kolmogorov–Smirnov tests.

**Supplementary Figure S5H:** Noise correlations were higher for intra-area than inter-area pairs among −VENs (*p* = 5.78 × 10⁻⁵³) and +VENs (*p* = 4.17 × 10⁻²⁹⁵), whereas correlations between −VEN/+VEN pairs did not differ between intra- and inter-area pairs (*p* = 0.137); Wilcoxon–Mann–Whitney test.

**Supplementary Figure S6B:** Areas enriched in neurons correlated with the tail vigour regressor (r ≥ 0.3) included ifDMO (*p* < 1 × 10⁻²⁵⁵), itVMO (*p* = 1.11 × 10⁻²⁰¹), Teg (*p* = 1.35 × 10⁻¹⁶⁷), NI (*p* = 5.02 × 10⁻¹³²), fMN (*p* = 5.48 × 10⁻¹¹³), nMLF (*p* = 6.69 × 10⁻¹¹²), PT (*p* = 1.41 × 10⁻⁷²), itDMO (*p* = 2.83 × 10⁻⁵⁹), sVMO (*p* = 3.44 × 10⁻⁴³), vMN (*p* = 1.15 × 10⁻²⁵), PTec (*p* = 1.90 × 10⁻²³), LC (*p* = 1.94 × 10⁻¹⁸), ifVMO (*p* = 1.29 × 10⁻¹⁶), VSL (*p* = 9.11 × 10⁻¹⁴), dTh (*p* = 5.88 × 10⁻¹³), Pal (*p* = 8.34 × 10⁻¹³), sDMO (*p* = 3.03 × 10⁻¹¹), POA (*p* = 5.05 × 10⁻⁶), HI (*p* = 2.12 × 10⁻³), SGN (*p* = 6.21 × 10⁻³), and SubP (*p* = 2.96 × 10⁻⁴); one-sided binomial tests performed per fish and combined across animals using Fisher’s method.

**Supplementary Figure S6C:** Areas enriched in neurons correlated with the straight bout regressor (r ≥ 0.3) included ifDMO (*p* < 1 × 10⁻²⁵⁵), itVMO (*p* = 4.57 × 10⁻⁹⁹), fMN (*p* = 3.07 × 10⁻⁴⁹), ifVMO (*p* = 4.43 × 10⁻³⁷), NI (*p* = 1.45 × 10⁻³⁹), sVMO (*p* = 3.18 × 10⁻¹⁸), Teg (*p* = 5.53 × 10⁻¹⁷), nMLF (*p* = 2.54 × 10⁻¹⁴), itDMO (*p* = 9.89 × 10⁻¹³), sDMO (*p* = 3.97 × 10⁻⁶), LC (*p* = 1.13 × 10⁻³), iRaphe (*p* = 5.86 × 10⁻³), vMN (*p* = 1.42 × 10⁻²), Cb (*p* = 2.21 × 10⁻²), and SGN (*p* = 4.22 × 10⁻²); one-sided binomial tests performed per fish and combined across animals using Fisher’s method.

**Supplementary Figure S7A:** Correlations with the tail vigour regressor were higher for −VENs than for +VENs (*p* = 1.47 × 10⁻¹⁶³); Wilcoxon–Mann–Whitney test.

**Supplementary Figure S7B:** Correlations of vigour-BCNs with the negative valence regressor were higher than correlations with the positive valence regressor (*p* < 1 × 10⁻²⁵⁵); Wilcoxon signed-rank test.

**Supplementary Figure S7D:** Stimulus-evoked noise correlations were greater than expected by chance for −VEN/vigour-BCN neuron pairs (*p* < 0.001) and +VEN/vigour-BCN neuron pairs (*p* < 0.001), comparison against matched-number randomly sampled neurons; Wilcoxon–Mann–Whitney test.

**Supplementary Figure S7E**: Spontaneous activity correlations were greater than expected by chance for −VEN/vigour-BCN neuron pairs (*p* < 0.001) and +VEN/vigour-BCN neuron pairs (*p* < 0.001), comparison against matched-number randomly sampled neurons; Wilcoxon–Mann–Whitney test.

**Supplementary Figure S7G**: Stimulus-evoked noise correlations for forebrain stimulus-specific neuron-turn-BCN pairs (Qun-turn-BCN, Cad-turn-BCN, Fex-turn-BCN, and Kin-turn-BCN) were significantly different from those of matched random pairs (*p* < 0.001 for all comparisons); Wilcoxon–Mann–Whitney test.

