## Supplementary Figures 1-7 for "Distributed representations of chemosensory valence in a naïve vertebrate brain"

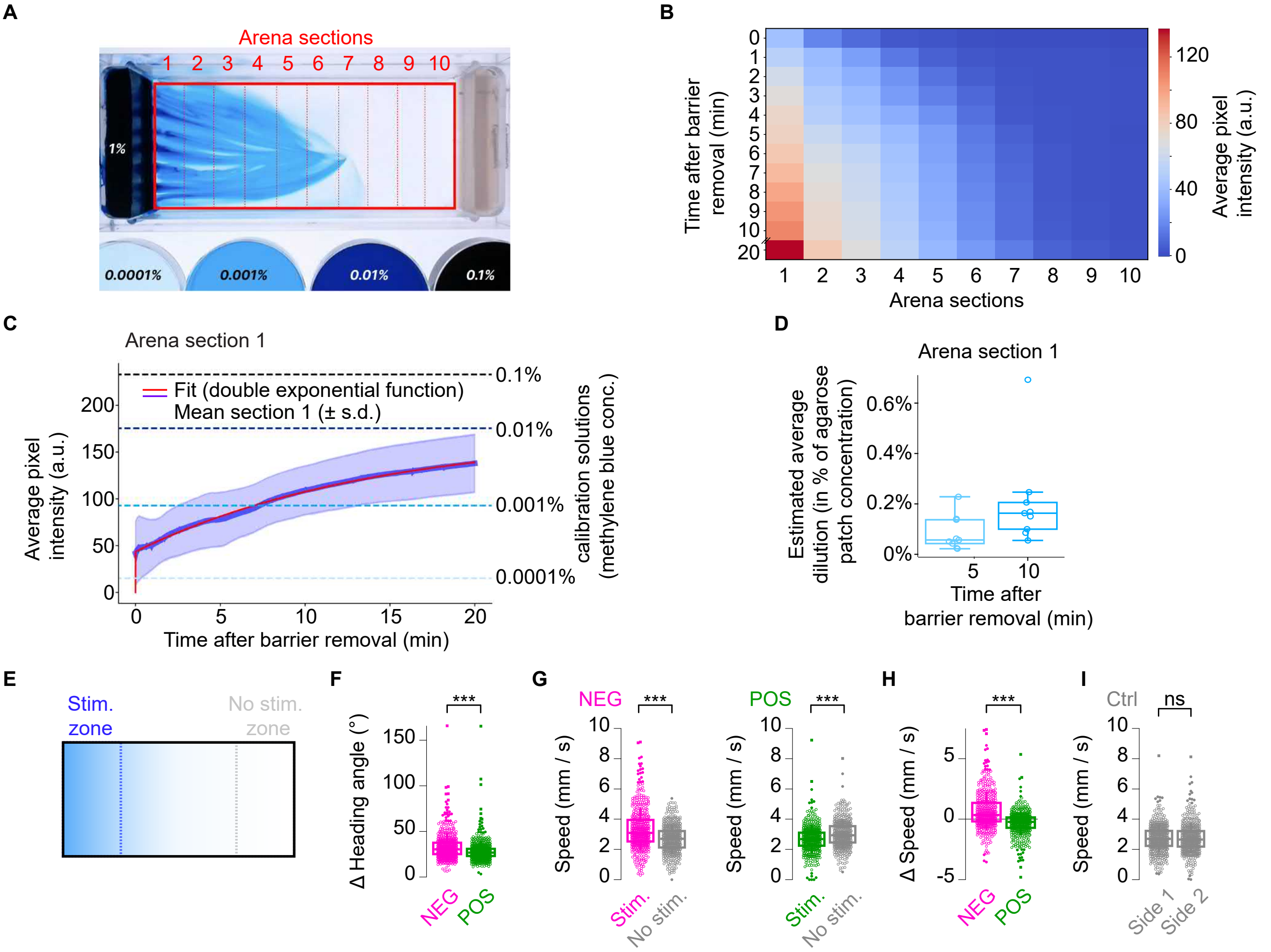


**Supplementary Figure S1. Estimated stimulus diffusion and single larvae analysis of behavioral responses to chemosensory stimuli in freely swimming naïve zebrafish larvae.** (**A**) Representative arena image 10 min after barrier removal, showing diffusion of methylene blue (1%) and calibration solutions with diluted dye. Red lines indicate arena sections used for analysis. (**B**) Heatmap of average pixel intensity across arena sections over time (N = 9 experiments). (**C**) Average pixel intensity over time in Section 1, closest to the dye source, compared with intensity values from calibration solutions. Shading indicates s.d.; red line shows a double-exponential fit. Right axis indicates pixel intensity values of the calibration solutions. (**D**) Methylene blue dilution in Section 1 at 5 and 10 min, estimated from calibration measurements and expressed as percentage of the source-patch concentration (1% methylene blue; N = 9 experiments). (**E**) Schematic showing the narrowly defined stimulus zone (Stim.) and the corresponding control zone near the agarose-only patch (No stim.). (**F**) Change in heading angle between consecutive bouts for aversive (NEG) and appetitive (POS) stimuli; Wilcoxon-Mann-Whitney test, $p = 8.6 \times 10^{-11}$; NEG, n = 527 fish; POS, n = 526 fish. (**G**) Swim speed within the stimulus zone compared with the corresponding no-stimulus zone for aversive stimuli (left; Wilcoxon signed-rank test, $p = 4.0 \times 10^{-15}$; n = 458 fish) and appetitive stimuli (right; $p = 2.1 \times 10^{-12}$; n = 443 fish). (**H**) Larva-wise swim-speed change relative to the no-stimulus zone for aversive and appetitive stimuli; Wilcoxon-Man-Whitney test, $p = 7.9 \times 10^{-8}$; n as in G. (**K**) Same analysis for agarose-only control experiments; Wilcoxon signed-rank test, p = 0.93; n = 464 fish.

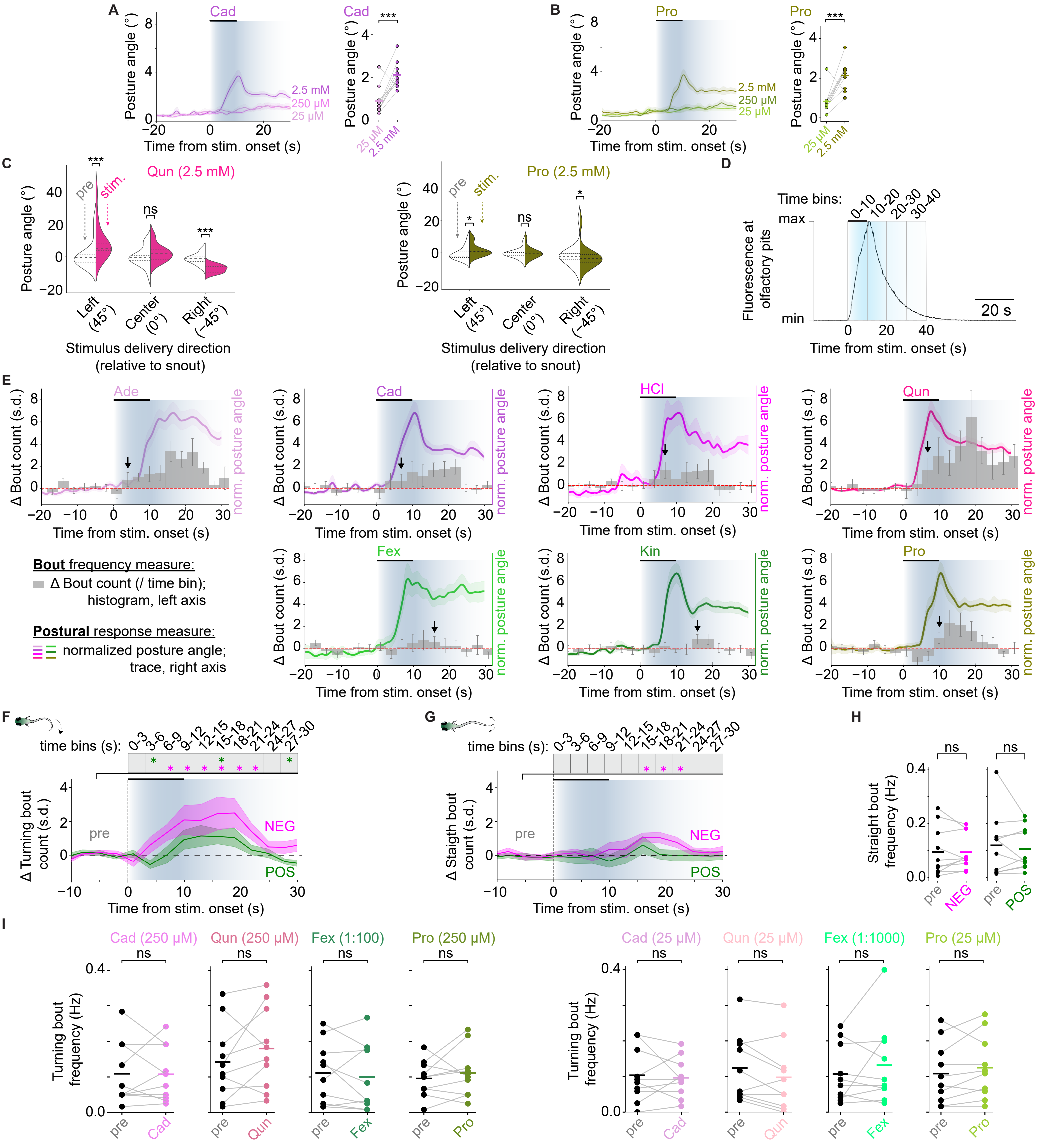


**Supplementary Figure S2. Additional analyses of behavioral responses to chemosensory stimuli in partially immobilized, naïve zebrafish larvae.** (**A**) Posture angle responses across three concentrations of Cad, shown as in **Fig. 2E**. The highest concentration elicited greater posture angle responses than the lowest concentration during the 20-s stimulus window; linear mixed-effects model with concentration as fixed effect, and fish identity (N = 10) and imaging session (forebrain or hindbrain) as random intercepts; ***$p < 0.001$. Background shading indicates approximate stimulus profile; trace shading indicates s.e.m. (**B**) Same as A, but for Pro. (**C**) Stimulus direction influences posture angle independently of stimulus valence. Violin plots show mean posture angle during the 20 s before and 20 s after chemosensory stimulus onset, with the stimulus delivery tube positioned 45° to the left, 45° to the right, or centered at 0° relative to the larva's snout. Left, Qun, 2.5 mM; right, Pro, 2.5 mM. Lateralized delivery elicited posture angle deflections toward the side of stimulus delivery for both stimuli. Linear mixed-effects model with time window (pre-stimulus vs. stimulus) as a fixed effect and fish identity (N = 5) as random intercept; ***$p < 0.05$, ***$p < 0.001$. (**D**) Peak-normalized fluorescein signal measured in a $40 \times 40\ \mu m^2$ region anterior to the olfactory epithelium, showing stimulus arrival and washout dynamics; representative measurement. The black bar indicates stimulus-channel activation; the peak occurred near the end of the 10-s stimulus window, followed by decay over the next ~20 s. (**E**) Peristimulus time histogram of pre-stimulus-normalized swim bout frequency, shown with peak-normalized mean posture angle responses to compare timing. Left axis: z-scored swim bout frequency (mean ± s.e.m.) over time relative to stimulus onset, binned in 3-s intervals, for each stimulus: Ade, 25 µM; Cad, 2.5 mM; HCl, pH 5; Qun, 2.5 mM; Fex, 1:10; Kin, 1:100; Pro, 2.5 mM. Shading indicates stimulus delivery and estimated washout; arrows mark the first bin exceeding the 95th percentile of baseline activity. Right axis: peak-normalized mean posture angle for the same stimuli; same data as in **Fig. 2D**. (**F**) Change in turning bout frequency over time, expressed as z-scored deviation from pre-stimulus baseline and quantified in 3-s bins, averaged across highest-concentration aversive (NEG; magenta) and appetitive (POS; green) stimuli; Wilcoxon signed-rank test versus baseline, *$p < 0.05$. Background shading indicates approximate stimulus profile; trace shading indicates s.e.m. (**G**) Same as F, but for straight bouts. (**H**) Straight bout frequency in 20-s pre-stimulus and stimulus windows for intermediate (left) and low (right) stimulus concentrations; mixed-effects model as in A, with stimulus condition as fixed effect. Each fish contributed six trials per stimulus-concentration pair. (**I**) Turning bout frequency in 20-s pre-stimulus and stimulus windows for individual stimuli at intermediate (left) and low (right) concentrations; analysis as in **Fig. 2J**; ns, not significant.

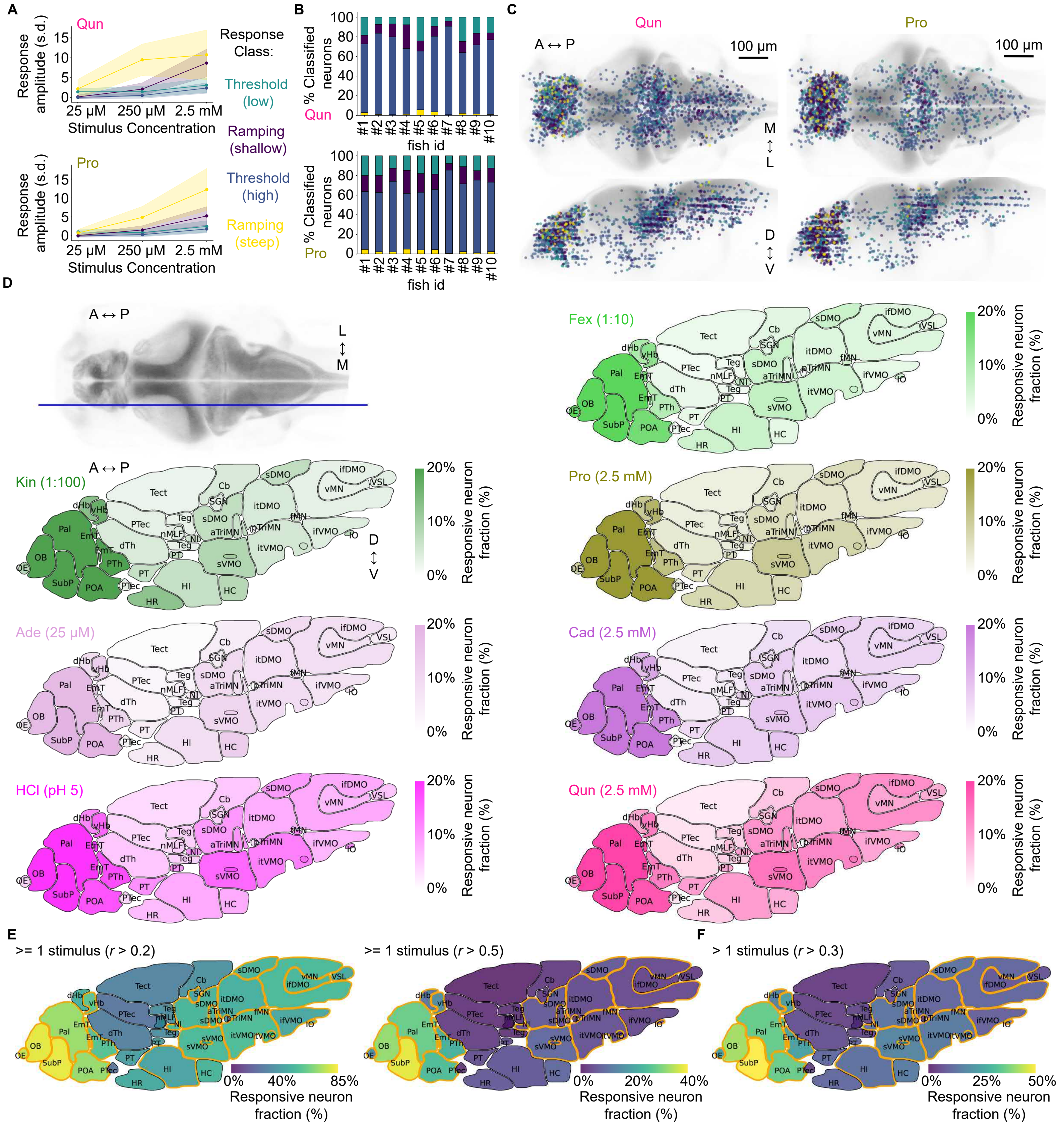


**Supplementary Figure S3. Additional analyses of chemosensory-evoked brain-wide neuronal activity.** (**A**) Average response amplitudes of neurons assigned to four concentration-response clusters for Qun (magenta, top) and Pro (ocher, bottom), shown as in **Fig. 3G**: cluster 1, steep-ramping (yellow); cluster 2, shallow-ramping (dark purple); cluster 3, low-threshold (turquoise); cluster 4, high-threshold (dark blue). (**B**) Proportions of each cluster type in forebrain and hindbrain FOVs for each stimulus and each fish. (**C**) Anatomical distribution of classified neurons registered to the reference brain and color-coded by cluster identity; 1 in 10 neurons shown for clarity. (**D**) Top left: reference slice used for mapping lateral brain regions. Remaining panels: fraction of neurons in each brain region with Pearson's $r > 0.3$ to the corresponding stimulus regressor, pooled across 10 fish. Stimuli: Fex, 1:10; Kin, 1:100; Pro, 2.5 mM; Ade, 25 µM; Cad, 2.5 mM; HCl, pH 5; Qun, 2.5 mM. Color indicates the fraction of responsive neurons per region. (**E**) Fraction of neurons in each brain region with Pearson's $r > 0.2$ (left) or $r > 0.5$ (right) to any highest-concentration stimulus regressor. (**F**) Fraction of neurons with Pearson's $r > 0.3$ to more than one highest-concentration stimulus regressor. Orange outlines indicate regions with significantly more responsive neurons than expected by chance; binomial test, FDR-corrected $p_{FDR} < 0.05$. A, anterior; P, posterior; D, dorsal; V, ventral; M, medial; L, lateral. Brain region abbreviations are defined in **Methods**.

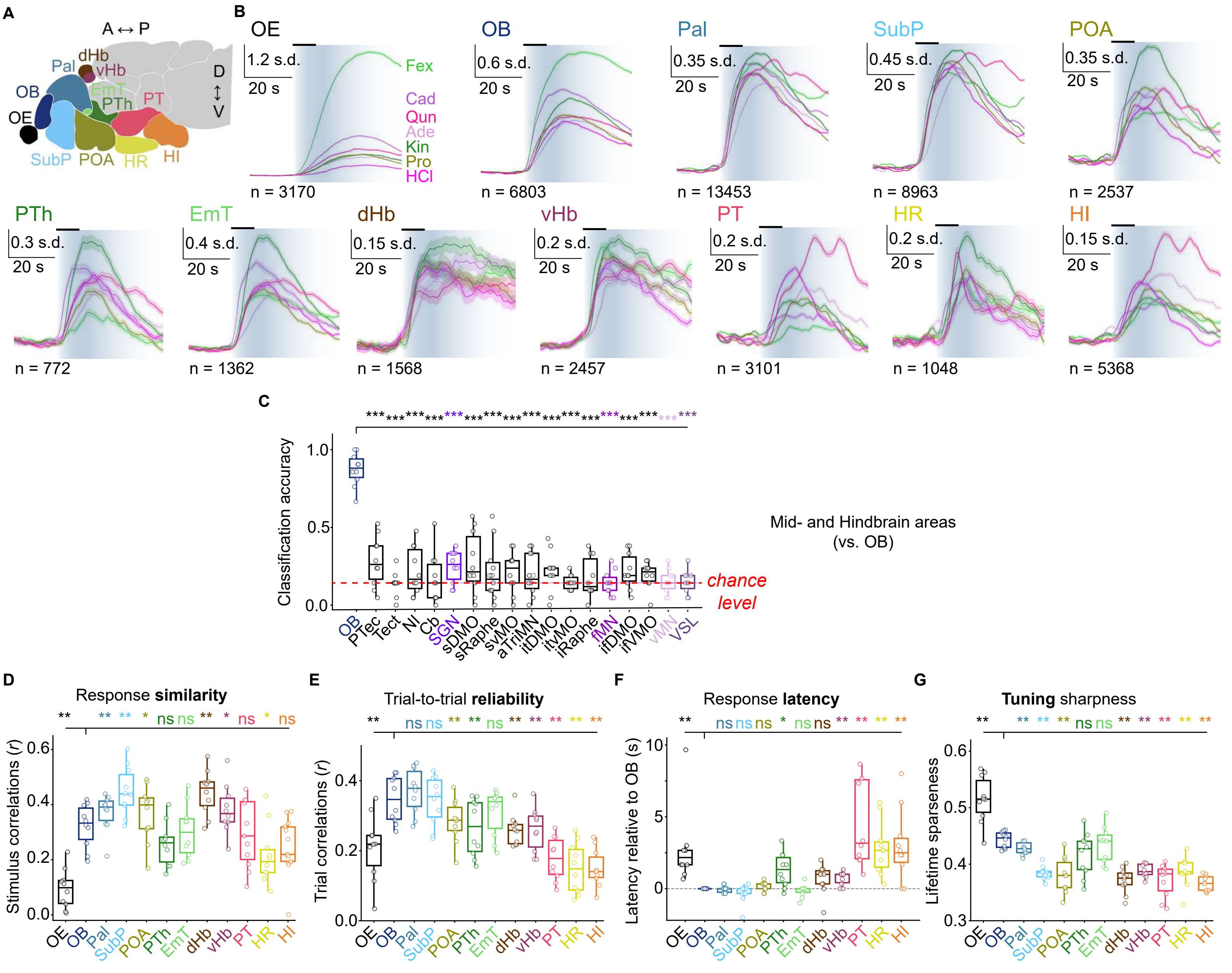


**Supplementary Figure S4. Region-specific response properties to chemosensory stimuli.** (**A**) Schematic sagittal section of the larval zebrafish forebrain, highlighting twelve highly responsive telencephalic and diencephalic regions. OE, olfactory epithelium; OB, olfactory bulb; Pal, pallium; SubP, subpallium; POA, preoptic area; PTh, prethalamus; EmT, eminentia thalami; dHb, dorsal habenula; vHb, ventral habenula; PT, posterior tuberculum; HR, rostral hypothalamus; HI, intermediate hypothalamus. (**B**) Trial-averaged, baseline-normalized z-scored $\Delta F/F(Ca^{2+})$ traces for each region in response to the seven highest-concentration stimuli. Lines show means and shaded envelopes indicate s.e.m. across neurons; n indicates number of neurons per region. Background shading indicates approximate stimulus profile. (**C**) Stimulus-classification accuracy across midbrain and hindbrain regions, using only highest-concentration stimuli. Asterisks indicate significant differences relative to OB; Wilcoxon signed-rank test, *p < 0.05, **p < 0.01, ***p < 0.001; ns, not significant. PTec, pretectum; TeO, optic tectum; NI, nucleus isthmi; Cb, cerebellum; SGN, secondary gustatory nucleus; sDMO, superior dorsal medulla oblongata; sRaphe, superior raphe; svMO, superior ventral medulla oblongata; aTriMN, anterior trigeminal motor nucleus; itDMO, intermediate dorsal medulla oblongata; itVMO, intermediate ventral medulla oblongata; iRaphe, inferior raphe; fMN, facial motor nucleus; ifDMO, inferior dorsal medulla oblongata; ifVMO, inferior ventral medulla oblongata; vMN, vagus motor nucleus; VSL, vagal sensory lobe. (**D**) Inter-stimulus pattern correlation, or response similarity, computed as the average Pearson correlation between population activity patterns evoked by different highest-concentration stimuli within each region. Wilcoxon signed-rank test, *p < 0.05, **p < 0.01, ***p < 0.001; ns, not significant; N = 10 fish. (**E**) Trial-to-trial correlation, or response reliability, computed as the average Pearson correlation between population responses to repeated presentations of the same stimulus (all stimulus-concentration conditions were considered). Asterisks indicate significant differences relative to OB; Wilcoxon signed-rank test, *p < 0.05, **p < 0.01; ns, not significant; N = 10 fish. (**F**) Global response latency after stimulus onset for each region, averaged across neurons and stimuli within each region and compared with OB. Wilcoxon signed-rank test, *p < 0.05, **p < 0.01, ***p < 0.001; ns, not significant; N = 10 fish. (**G**) Lifetime sparseness, or 'tuning sharpness', of neuronal responses across regions. Individual data points show region-specific fish averages. Asterisks indicate significant differences relative to OB; Wilcoxon signed-rank test, *p < 0.05, **p < 0.01, ***p < 0.001; ns, not significant; N = 10 fish.

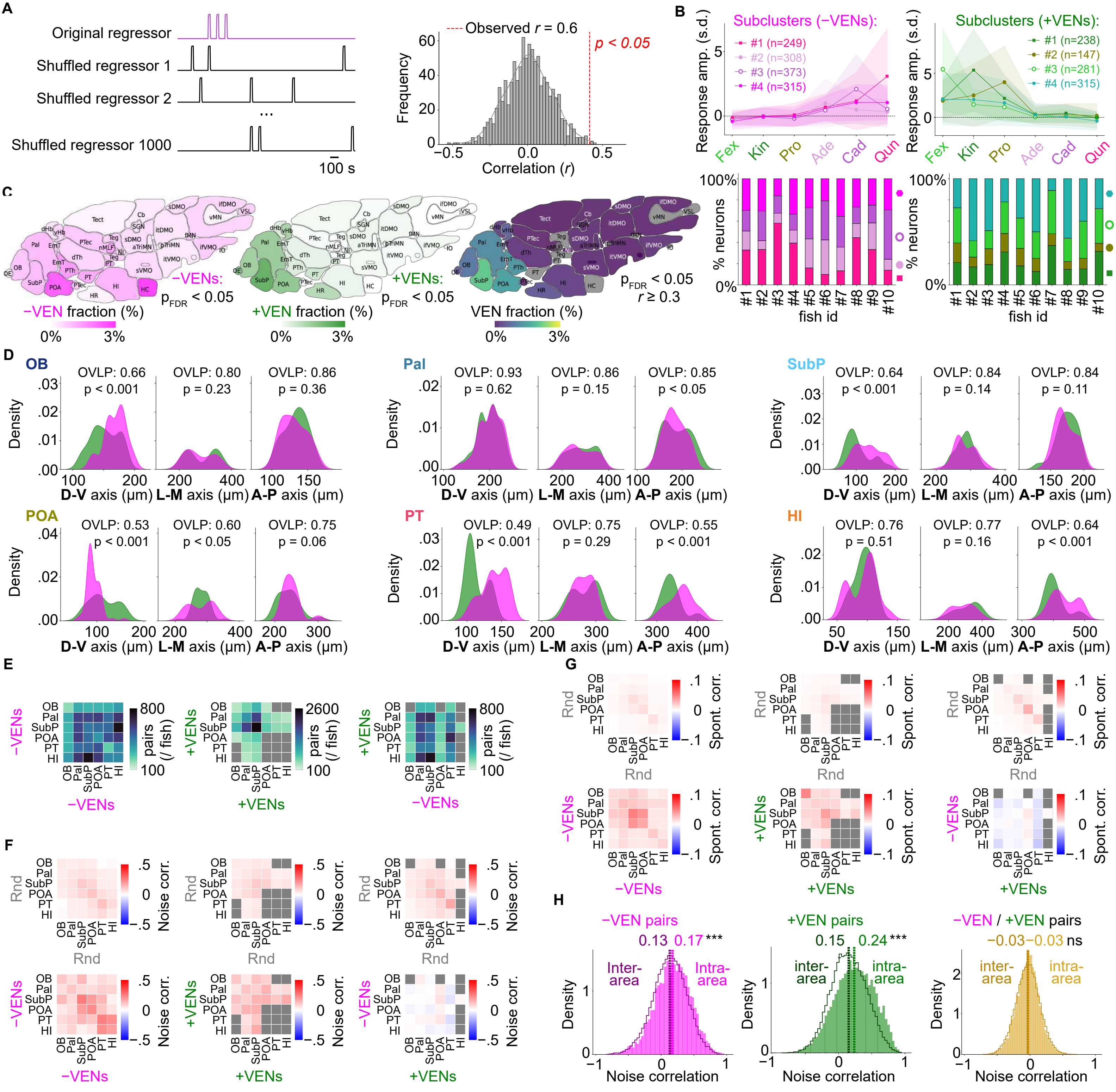


**Supplementary Figure S5. Additional analyses of valence-encoding neurons (VENs) and their functional coupling.** (**A**) Schematic of permutation-based significance testing for neuron-regressor correlations. Left, example original stimulus or behavior regressor (top, purple) and randomly shuffled regressors (black) used to generate a null distribution. Shuffling disrupts temporal event alignment while preserving event number and shape. Right, example null distribution of correlation values from 1,000 shuffled regressors. The observed correlation for one neuron (Pearson's $r = 0.6$; dashed red line) lies outside the 95th percentile of the null distribution, indicating a significant correlation ($p < 0.05$). (**B**) Top: mean ± s.d. response profiles of −VENs (left) and +VENs (right) grouped by k-means clustering; n indicates number of neurons per cluster. Bottom: distribution of VEN clusters across individual fish. Stacked bars show the fraction of +VENs (left) and −VENs (right) assigned to each k-means cluster (k = 4), grouped by fish; colors and symbols indicate cluster identity. Similar cluster proportions across fish indicate that these VEN response-profile classes were reproducible across fish. (**C**) Spatial distribution of VENs across brain regions. Left: fraction of significant −VENs ($p_{FDR} < 0.05$). Middle: fraction of significant +VENs ($p_{FDR} < 0.05$). Right: fraction of VENs ($p_{FDR} < 0.05$) exceeding an additional correlation threshold (Pearson's $r > 0.3$). Color saturation indicates the fraction of VENs relative to all neurons in each region, aggregated across fish. (**D**) Spatial gradients of VENs along principal anatomical axes. Distributions of +VENs (green) and −VENs (magenta) are shown along the dorsoventral (D-V), mediolateral (M-L), and anterior-posterior (A-P) axes in six VEN-rich brain regions. Overlap scores (OVLP) and Kolmogorov-Smirnov test p-values are shown for each axis and region. (**E**) Average VEN-pair counts per fish for each region pair. Gray squares indicate region pairs with <100 average neuron pairs, which were excluded from further analysis. (**F**) Top: mean noise correlations for number-matched random neuron pairs within and across six forebrain VEN-rich regions. Bottom: mean noise correlations for VEN pairs within and across the same regions. Data related to **Fig. 5J**. (**G**) Same as F, but for spontaneous correlations. Data related to **Fig. 5M**. (H) Noise correlation distributions for intra- and inter-area −VEN/−VEN pairs (n = 12,682 and 27,812), +VEN/+VEN pairs (n = 29,626 and 32,217), and −VEN/+VEN pairs (n = 8,731 and 33,993). OB, olfactory bulb; Pal, pallium; SubP, subpallium; POA, preoptic area; PT, posterior tuberculum; HI, intermediate hypothalamus. Other abbreviations are defined in **Methods**. A, anterior; P, posterior; D, dorsal; V, ventral; M, medial; L, lateral.

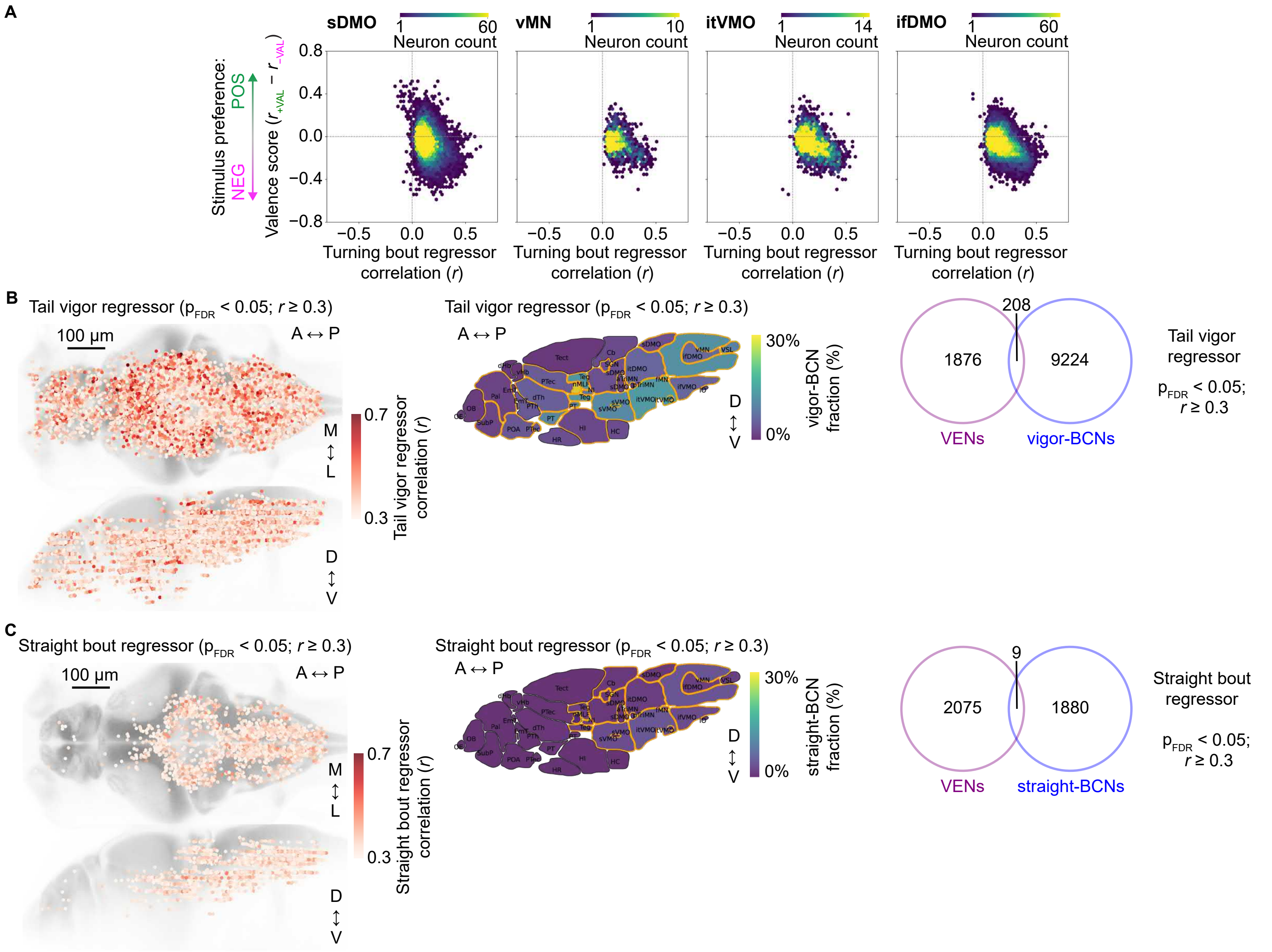


**Supplementary Figure S6. Additional analyses of behaviorally correlated neurons (BCNs).** (**A**) Joint distribution of valence score and turning bout regressor correlation across VENs and turn-BCNs in selected hindbrain regions. Each dot represents one neuron, plotted by its valence score, defined as *r*(+VAL) − *r*(−VAL), and its Pearson correlation *r* with the turning bout regressor. Color indicates local neuron density. sDMO, superior dorsal medulla oblongata; vMN, vagus motor nucleus; itVMO, intermediate ventral medulla oblongata; ifDMO, inferior dorsal medulla oblongata. (**B**) Spatial distribution and regional enrichment of vigor-BCNs, defined as neurons significantly and strongly positively correlated with the tail vigor regressor ($p_{FDR} < 0.05$; $r \geq 0.3$). Left: spatial distribution of vigor-BCNs. Middle: fraction of vigor-BCNs across brain regions, pooled across fish; orange outlines indicate significant regional enrichment; binomial test, FDR-corrected $p_{FDR} < 0.05$. Right: overlap between VENs ($p_{FDR} < 0.05$) and vigor-BCNs. In total, 208 neurons were classified as both VENs and vigor-BCNs, corresponding to ~10% of VENs and ~2% of vigor-BCNs. (**C**) Same as B, but for straight-BCNs identified using the straight bout regressor. A, anterior; P, posterior; D, dorsal; V, ventral; M, medial; L, lateral. Additional brain region abbreviations are defined in **Methods**.

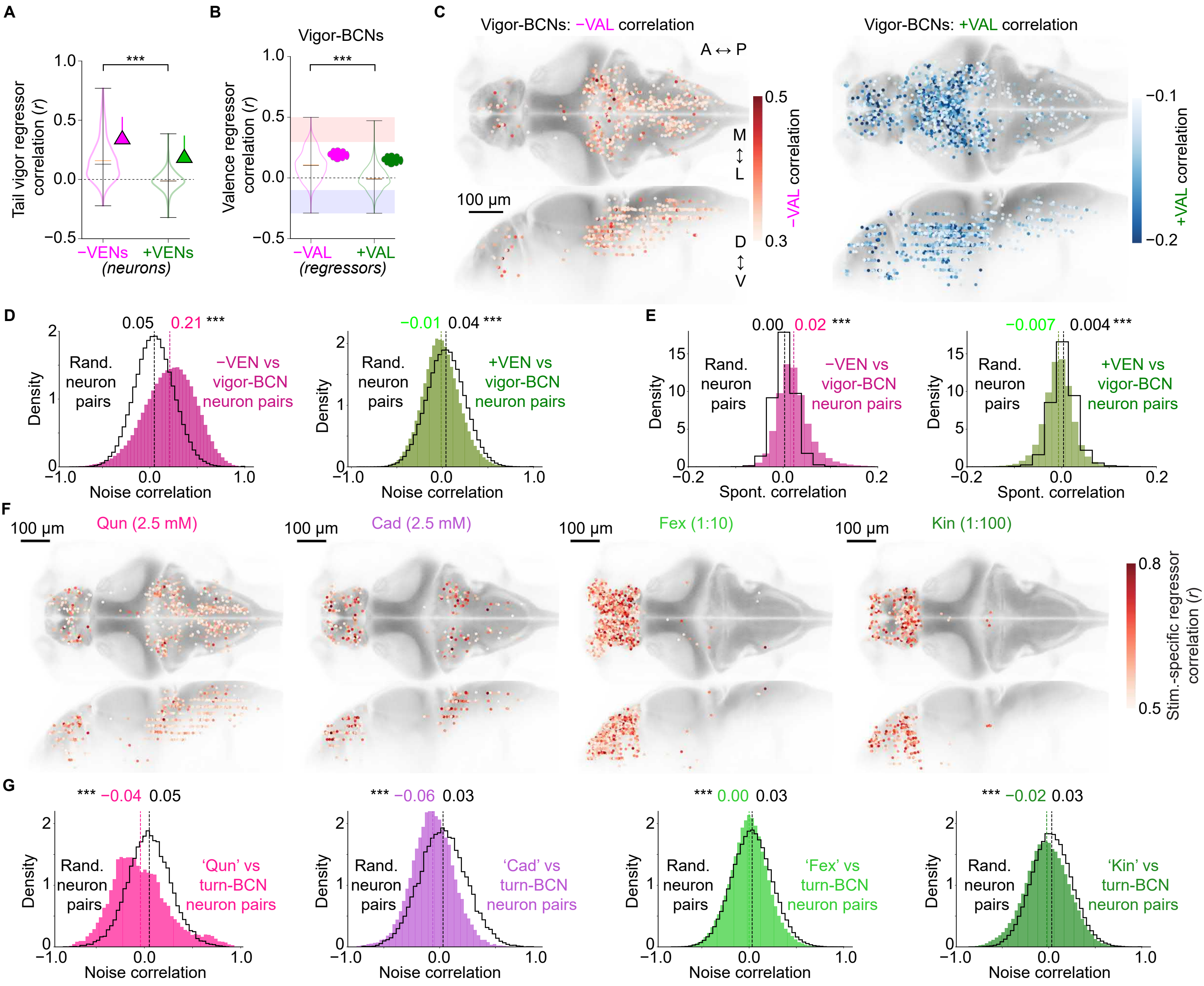


**Supplementary Figure S7. Additional analyses of sign-specific relationships between VENs and behavior-correlated activity.** (**A**) Correlations of −VENs and +VENs with the tail vigor regressor. Black line, median; orange line, mean; Wilcoxon-Mann-Whitney test, ***p < 0.001. (**B**) Correlations of vigor-BCNs with −VAL and +VAL valence regressors; Wilcoxon-Mann-Whitney test, ***p < 0.001. (**C**) Anatomical distributions of vigor-BCNs positively correlated with −VAL ($r \geq 0.3$; left, warm colors) or negatively correlated with +VAL ($r \leq -0.1$; right, cool colors). (**D**) Noise correlation distributions for −VEN-vigor-BCN pairs (magenta; n = 265,034) and +VEN-vigor-BCN pairs (green; n = 199,119), compared with matched random pairs (black). (**E**) Same as D, but for spontaneous correlations. (**F**) Brain-wide maps of stimulus-specific neuronal responses. Each panel shows neurons significantly and strongly correlated with a stimulus-specific regressor ($p_{FDR} < 0.05$; $r \geq 0.5$) for one chemosensory stimulus: Fex (n = 1092), Kin (n = 325), Cad (n = 215), and Qun (n = 504). Dorsal (top) and lateral (bottom) views are shown for each stimulus. Each dot represents one neuron, colored by Pearson's *r* with the corresponding stimulus regressor. (**G**) Noise correlation distributions for forebrain stimulus-specific neuron-turn-BCN pairs (Qun-turn-BCN, n = 13,569 pairs; Cad-turn-BCN, n = 20,589; Fex-turn-BCN, n = 205,983; Kin-turn-BCN, n = 116,638), compared with matched random pairs (black).
